# Unexpected changes in reproductive barriers between incipient species after experimental evolution in sympatry

**DOI:** 10.64898/2026.02.06.704315

**Authors:** Miguel Cunha, Miguel A. Cruz, Inês Santos, Vitor C. Sousa, Sara Magalhães, Leonor R. Rodrigues, Flore Zélé

## Abstract

Hybridization is generally considered a temporary phenomenon, but it is actually widespread and may last for large time periods between species that stably coexist. Here, to test whether evolving with a closely-related species modifies or maintains partial reproductive isolation, we performed experimental evolution in artificial sympatry *vs*. allopatry with two closely-related colour forms of spider mites (*Tetranychus urticae*) that exhibit an asymmetrical pattern of pre-mating isolation despite almost complete post-zygotic isolation. We assessed whether evolutionary changes occurred in traits associated to (i) pre-mating isolation, (ii) post-mating pre-zygotic and early post-zygotic isolation, and (iii) late post-zygotic isolation. Our results revealed that reinforcement did not occur even under forced long-term sympatric evolution. Instead, the strength of some reproductive barriers decreased (*e.g*., premating isolation and fertilization failure), and some trait changes indicated convergence rather than divergence between species (*e.g*., mating propensity, latency to copulation). In fact, both types of males showed the same decreased preference for red-form females across generations in sympatry. In line with this, traits underlying fertilization success evolved in the same direction and with similar amplitude in heterotypic crosses and in their homotypic control, as the offspring sex ratio of green-form females decreased in sympatry irrespective of the male they mated with. Finally, other changes in reproductive barriers resulted from trait correlations (*e.g*., decreased zygote mortality but increased juvenile mortality). Hence, despite very high costs of hybridization, responses occurring following evolution in sympatry were unrelated to selection directly associated to hybridization, but rather the by-product of other evolutionary forces, with cascading consequences for reproductive barriers. In particular, these results support the underappreciated hypothesis that within-species sexual interactions can constrain population divergence, or even drive trait convergence between species, thereby playing a role in the maintenance of partial reproductive isolation.

## Introduction

Despite decades of research on speciation, we still lack a clear understanding of the processes driving the evolution of reproductive barriers, eventually leading to the origin of reproductively isolated species. Patterns of naturally occurring reproductive isolation provided great insight into the mechanisms underlying its evolution (Coyne & Orr, 1989, 1997; Matute & Cooper, 2021). For instance, field studies showed that prezygotic isolation is often stronger between sympatric than between allopatric taxa (*e*.*g*., Sætre *et al*., 1997; Yukilevich, 2012; Nosil, 2013), and pairs of taxa with partially overlapping geographic ranges frequently show enhanced isolation in sympatry (Gerhardt, 1994; Noor, 1995; Rundle & Schluter, 1998; Matute, 2010, reviewed in Shaw *et al*., 2024). These patterns are generally interpreted as the product of reinforcement, whereby prezygotic isolation between divergent taxa is strengthened during secondary contact due to selection arising from reduced hybrid fitness (Dobzhansky, 1940; Marshall *et al*., 2002; Servedio, 2004), or, more generally, as the result of reproductive character displacement driven by reproductive interference (Brown & Wilson, 1956; Blair, 1974; Kyogoku, 2015).

Reproductive interference refers to the negative effect that interspecific reproductive sexual interactions, such as signal jamming, misdirected mating courtship, mating attempts, successful copulations, and hybridization, have on the fitness of the species involved (Gröning & Hochkirch, 2008; Burdfield-Steel & Shuker, 2011). It has been reported in organisms of many different taxa, ranging from angiosperms to vertebrates and has received much attention from evolutionary biologists for its effect on speciation (reviewed in Gröning & Hochkirch, 2008; Kyogoku, 2015). Moreover, it has gained growing recognition as a major driver of species exclusion (Kishi *et al*., 2009; Schreiber *et al*., 2019; Yamamichi *et al*., 2023; Cruz *et al*., 2025a). Therefore, patterns of character displacement observed in sympatry may simply be due to the fact that taxa with stronger prezygotic isolation (hence lower reproductive interference) are more likely to persist in sympatry than are weakly isolated ones (which either fuse or go extinct; Templeton, 1981; Weber & Strauss, 2016; Anderson & Matute, 2025).

Alternatively, trait changes in species that co-occur in sympatry may result from other selective forces (Maan & Seehausen, 2011). For example, different types of interactions, such as predation (Jiggins *et al*., 2001; Arias *et al*., 2008) or competition for shared resources (Day, 2000; Bürger *et al*., 2006; Winkelmann *et al*., 2014) can shape reproductive isolation if the ecological traits under selection also impact reproductive interactions (Dodd, 1989; Rice & Salt, 1990; Nagel & Schluter, 1998; Pfennig & Pfennig, 2009; Hoskin & Higgie, 2010). In addition, sexual selection and sexual conflict operating within species have been identified as powerful engines of speciation (Ritchie, 2007; Gavrilets, 2014, but see Kraaijeveld *et al*., 2011). Yet, both natural and sexual selection may also have opposite effects, hampering divergence or even driving trait convergence (*e*.*g*., Drury *et al*., 2015; Llaurens *et al*., 2021; Simpson *et al*., 2021; Puissant *et al*., 2023). This may explain the widespread occurrence of partial reproductive isolation in natural populations, despite high costs of reproductive interference (Cotto & Servedio, 2017; Taylor *et al*., 2017; Servedio & Hermisson, 2020).

Although comparative studies can elegantly hint at mechanisms underlying a speciation event (*e*.*g*., Rundle & Schluter, 1998; Hopkins & Rausher, 2012; Yukilevich, 2012; Nosil, 2013), tracking the evolutionary responses of replicated populations exposed to manipulated controlled conditions (*i*.*e*., experimental evolution) remains one of the most robust approaches to identify such mechanisms (White *et al*., 2020). These studies have demonstrated the role of adaptation to different environments, while depreciating the role of population bottlenecks and of disruptive selection as main drivers of speciation (*e*.*g*., Rice & Hostert, 1993; Fry, 2009; Matute, 2010a, 2015; Myers & Frankino, 2012; White *et al*., 2020). They also showed that prezygotic isolation can evolve in sympatry when all F1 hybrids produced between two populations are removed at each generation (Fry, 2009; Coughlan & Matute, 2020; Jezovit & Levine, 2025). However, very few experimental evolution studies have tested the consequences of secondary contact in which hybrids are not killed by design, which is highly relevant to understand patterns of reproductive isolation observed in natural populations (Higgie *et al*., 2000; Coughlan & Matute, 2020; Matute & Cooper, 2021; Jarvis *et al*., 2024). Here, we used such an evolution experiment to shed light on an existing pattern of asymmetrical premating isolation (*i*.*e*., both assortative and disassortative mating), despite strong intrinsic postzygotic barriers, between divergent populations (or subspecies) naturally occurring in the same geographic area.

In spider mites, and particularly within the genus *Tetranychus* (Acari: Tetranychidae), reproductive barriers are often incomplete between species and variable degrees of reproductive isolation can be found within species (*e*.*g*., Clemente *et al*., 2016; Cruz *et al*., 2021; Xue *et al*., 2023; Cruz *et al*., 2025b). This provides an excellent opportunity to study their evolution at different points in the speciation continuum (Kulmuni *et al*., 2020). The short generation time of these phytophagous arthropods (10-16 days at 25ºC; Riahi *et al*., 2013) allows performing experimental evolution within a relatively short timeframe (Magalhães *et al*., 2007a; Bitume *et al*., 2011; Rodrigues *et al*., 2022; Fragata *et al*., 2025), and their haplodiploid sex determination facilitates investigations of fertilization failure, because sons develop from unfertilized eggs and daughters from fertilized eggs (*i*.*e*., true arrhenotoky; Helle & Bolland, 1967; Oku, 2014; Clemente *et al*., 2018; Cruz *et al*., 2021, 2025b). These mites also exhibit sexual dimorphism and their virginity can be ensured by isolating individuals just before the last moulting stage. Finally, females mated with conspecific (or homotypic) males exhibit first-male sperm precedence, whereby the sperm of the first male mating with a female is used to fertilize most eggs (Helle, 1967; Rodrigues *et al*., 2020). Thus, if this pattern is maintained when a female mates with a male from a strongly divergent population or semi-isolated species, the costs of reproductive interference should be particularly high, and thus constitute a strong selective pressure for the evolution of reproductive barriers by reinforcement (Shuker & Burdfield-Steel, 2017).

Here, we used a pair of populations from genetically differentiated colour forms of the spider mite *Tetranychus urticae*: the green and red forms, which have worldwide overlapping distributions and host plant ranges (Migeon & Dorkeld, 2025), and can co-occur on the same individual host plant (Lu *et al*., 2017; Zélé *et al*., 2018). Due to complete reproductive isolation among some populations of these two forms, they have been described as separate species: *T. urticae* and *T. cinnabarinus*, respectively (*e*.*g*., Smith, 1975b). However, because other populations produce viable hybrids over multiple generations (*e*.*g*., Gotoh & Tokioka, 1996; Sugasawa *et al*., 2002), and morphological and molecular data suggests synonymity (Auger *et al*., 2013, but see Xue *et al*., 2023 for more recent genomic data), they are currently considered two ‘forms’ or ‘incipient species’, with variable degrees of reproductive incompatibilities (Dupont, 1979; De Boer, 1982; Sugasawa *et al*., 2002; Cruz *et al*., 2021; Xue *et al*., 2023). In particular, previous laboratory studies performed with the two populations used in the present study revealed a strong male-biased sex ratio distortion in heterospecific crosses between green-form females and red-form males (Cruz *et al*., 2021). Furthermore, both directions of heterotypic crosses result in fully sterile hybrids (most hybrid females do not lay eggs and the few eggs laid do not hatch; Cruz *et al*., 2021). Yet, irrespective of their colour form, females do not display mate preference, and males of both colour forms prefer to mate with red-form females (Cruz *et al*., 2025b). Such incomplete and asymmetrical pre-zygotic isolation (Cruz *et al*., 2025b), despite strong post-zygotic isolation (Cruz *et al*., 2021), is puzzling, especially given first male sperm precedence (Helle, 1967; Rodrigues *et al*., 2020). Yet, this pattern may just be due to limited contact between the two populations in the field, before being brought to the laboratory, even though they were collected in the same region (see details in Box S1). Alternatively, secondary contacts between populations of the two forms may be frequent enough, but reinforcement may be prevented by other evolutionary forces (Ritchie, 2007; Servedio & Hermisson, 2020). To test this, while limiting the role of ecological speciation (Schluter, 2001), we performed experimental evolution of these populations either together in artificial sympatry, or separately in allopatric conditions for up to 69 generations in a homogeneous environment. We then measured the strength of all reproductive barriers between evolved populations: (1) pre-mating isolation (mate choice and genitalia incompatibilities), (2) post-mating pre-zygotic isolation (barriers to sperm transfer/storage, gametic incompatibilities and homotypic sperm precedence) and early post-zygotic isolation (zygote mortality and hybrid inviability); and (3) late post-zygotic isolation (hybrid sterility and hybrid breakdown).

## Materials and methods

### Spider mite source populations

Two spider mite source populations, belonging to the green and red form of *T. urticae* and fully described in supplementary Box S1, were used in this study. Briefly, these populations were collected in nearby locations in central Portugal, treated with antibiotics to obtain symbiont-free populations, then reared in mite-proof cages containing bean plants (*Phaseolus vulgaris*, cv. Contender seedlings obtained from Germisem, Oliveira do Hospital, Portugal), under standard laboratory conditions (*ca*. 500-1000 females; 24 ± 2ºC, 16/8h L/D), before starting experimental evolution.

### Experimental evolution

Three selection regimes were created in December 2018 using the green- and red-form symbiont-free source populations described above (Figure 1): green-form or red-form individuals evolving in absence of heterotypic individuals (“allopatry” regimes) and individuals from the two forms evolving together (“sympatry” regime), each composed of five independent population replicates. Each replicate population was created by transferring 200 young adult mated females (either from a single species – allopatry regimes; or from both species – sympatry regime) from the source populations to plastic boxes containing two 14-days-old bean plants. A fresh bean plant was added to each experimental box 7 days later to avoid resource depletion, and the same number of mated female offspring were transferred to new boxes with two fresh plants at each generation (every 14 days). To prevent the loss of the red-form populations in the sympatry regime, we then increased the number of red-form females transferred from 200 to 400 per replicate population of this regime starting in July 2019 (T17). Still, a replicate population was lost in December 2020. To prevent further losses from January 2021 (T55) onwards, back-up populations were created at each generation with 200 to 400 red-form mated female offspring, to be used if not enough red females were found in the sympatry replicate populations (see the detailed procedure in box S2). The experimental evolution, as well as all common gardens and experiments described below were all conducted in a growth chamber with standard conditions (24 ± 2ºC, 60% RH, 16/8h L/D).

**Figure 1.**
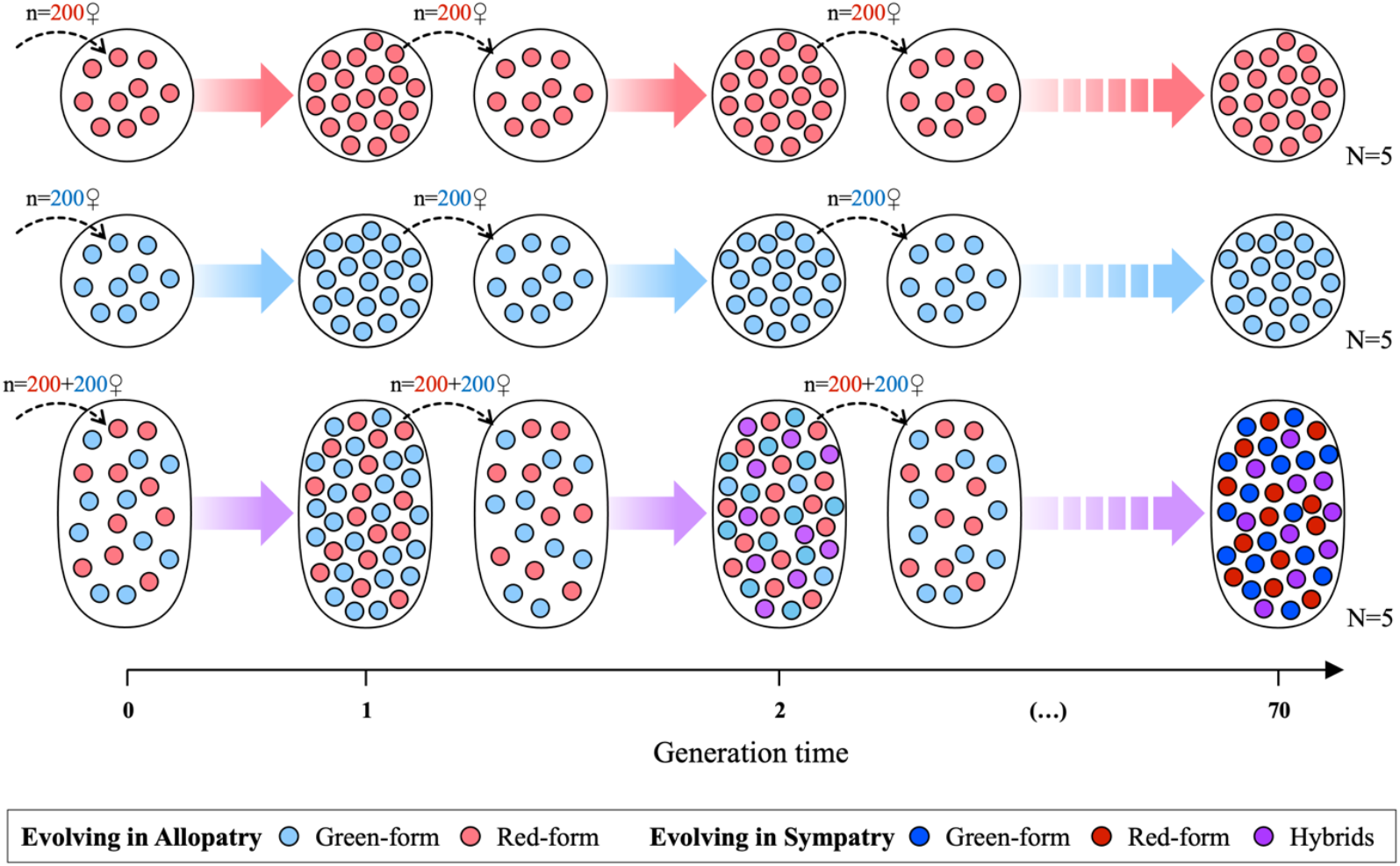
Overview of the procedure used for experimental evolution of red- and green-form spider mites in artificial sympatry *vs* allopatry. Five independent population replicates of each *T. urticae* colour form evolved separately or together (allopatry and sympatry regimes, respectively) for up to 70 generations. Coloured arrows indicate offspring production and development whereas black dashed arrows indicate the transfer of mated females to a new experimental box at each generation (see box S2 for more details).

### Common gardens

Common gardens were performed for green- and red-form females of each selection regime and replicate population prior to each experiment. In addition to reducing the influence of environmental/maternal effects (Kawecki & Ebert, 2004), this procedure ensured that all males used in experiments were of the correct colour form, and that none of the red females originating from the sympatry regime were F1 hybrids, given that both red-form and hybrid females are red (see box S1). Two generations before the onset of an experiment, 100-200 mated females (depending on the experiment) of each colour form and from each population replicate of each selection regime, were placed separately in a new plastic box containing two fresh bean plants. To avoid resource depletion, a fresh bean plant was subsequently added to each box ca. every 7 days. Then, mated female offspring were collected from each box 10 to 23 days after the establishment of the common gardens to create the ‘age-cohorts’ described below for each experiment.

### Pre-mating reproductive barriers: mate preference and genitalia incompatibilities (Experiment 1)

To test for the evolution of pre-mating reproductive barriers between populations of the two colour-forms, mate preference tests were conducted following 50 and 67-69 experimental evolution transfers (‘T50’ and ‘T67-69’ hereafter; see box S2 for corresponding effective generations of selection). Only male mate preference was tested (Figure 2A) because opportunities for females to choose a partner in this system are expected to be very low (hence the window for selection to act on this trait), as males usually guard quiescent deutonymph females (teleiochrysalis) and mate with them as soon as they emerge as adults (Oku 2014). In line with this, a previous study with these populations showed that females of the two colour forms do not prefer males of either form, whereas males of the two colour forms prefer red-form females (Cruz *et al*., 2025b).

**Figure 2.**
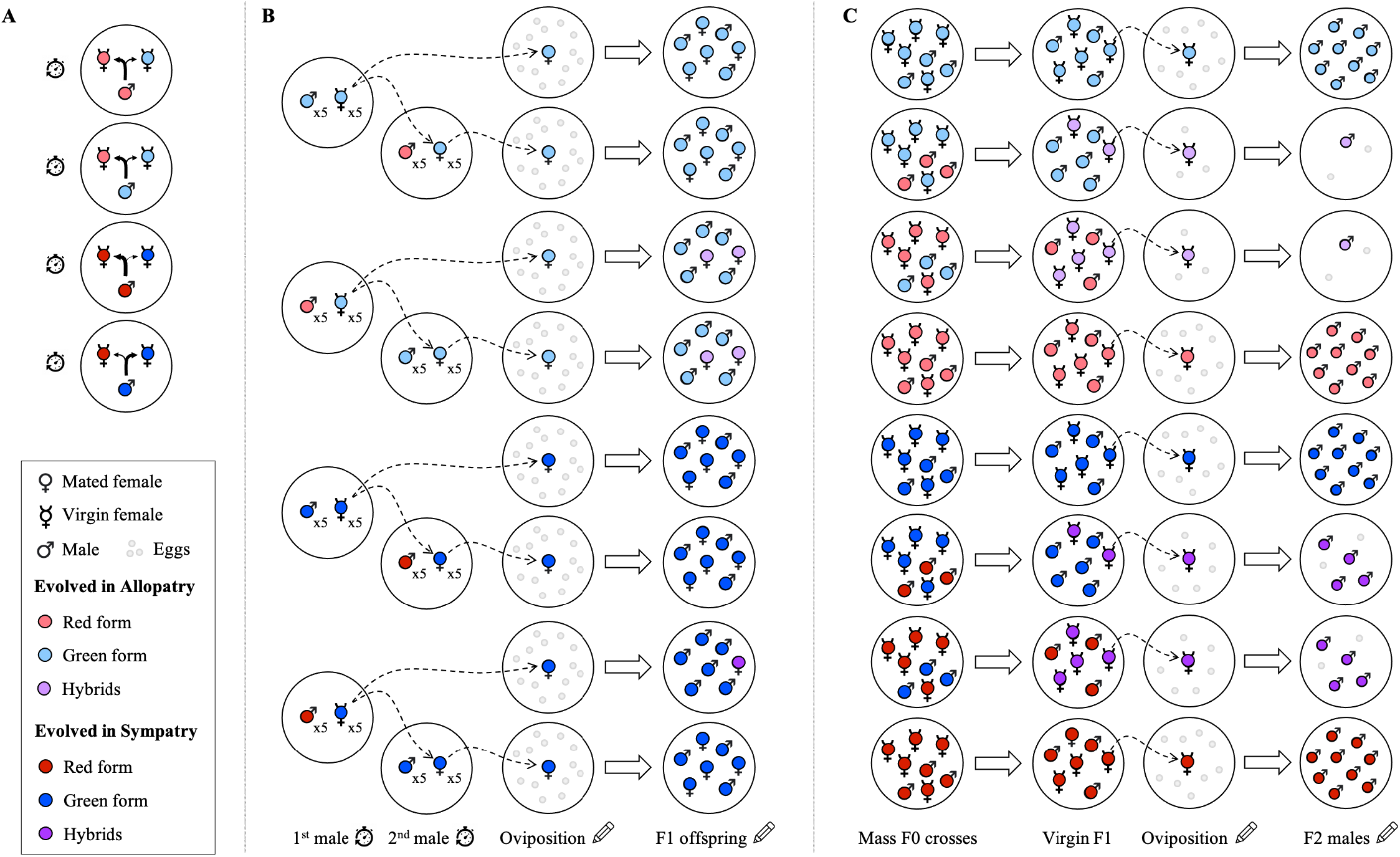
Overview of experimental procedures and hypothetical scenarios of evolution of different reproductive barriers: A - mating preference (Experiment 1), B - fertilization failure, sperm precedence & F1 hybrid mortality (Experiment 2), and C - F1 hybrid fertility and F2 hybrid breakdown (Experiment 3), between the green and red-form populations evolving in sympatry *vs* allopatry. Depicted scenarios correspond to reinforcement of pre-zygotic isolation, but decreased post-zygotic isolation due to gene-flow, after evolution in sympatry, with the evolution of homotypic male preferences in both cross directions (A), increased fertilization failure and evolution of homotypic sperm precedence instead of first male sperm precedence (B), and increased F1 hybrid fertility and decreased F2 hybrid breakdown (C). In (A), thicker arrows indicate a stronger preference. In (B) and (C), dashed thin arrows indicate the transfer of a female from one leaf disc to another, and large white arrows indicate offspring production and development. Clock and pencil icons indicate variables that were measured (time, eggs and adult offspring).

To obtain males and virgin females roughly of the same age for the preference tests, age-cohorts were created 11-12 days before mating observations, by transferring mated females from each common garden onto a bean leaf placed on water-soaked cotton (50 females from the allopatric regime and 80-100 from the sympatry regime to compensate for the risk of transferring hybrid females). Females were allowed to lay eggs either for two days on the same bean leaf then transferred to a new fresh leaf for another two days (at T50), or for three days on the same bean leaf (at T67-69). Females and males were then isolated from each age-cohort as soon as they reached their last teleiochrysalis stage. Each sex was kept separately on 2.5 cm^2^ leaf discs to ensure their virginity when emerging as adult. Mate preference tests were then performed either the next day (T50) or two days later (T67-69).

For each preference test, a green-or red-form male, originating from the allopatry or sympatry regime, was given the choice between a green- and a red-form female from its own selection regime (and from its own population replicate, or equivalent in the case of the allopatry regime. *E*.*g*., males from replicate 1 were always exposed to green- and red-form females of replicate 1). The two females of each test were installed on a 0.5 cm^2^ leaf disc (hereafter called ‘arena’), followed by the focal male. Time started counting when the male was added to the arena, with the seconds the copulation started and finished, as well as the colour of the female chosen by the focal male, being noted by the observer. If no copulation occurred within 30 minutes, the arenas were excluded from the statistical analyses. Timings were recorded with a stopwatch (Microsoft Application “Stopwatch: StopNow” v. 3.0.1, by Stefan Spittank) and used to calculate latency to copulation (time between male placement on the disc and beginning of copulation, in seconds) and copulation duration (time in seconds males spent copulating).

Different sessions of observation were performed, each allowing to test for the preference of either two (at T50) or three (at T67-69) males of each colour form (green *vs*. red), selection regime (allopatry *vs*. sympatry) and population replicate (N=4 for this experiment), hence a total of 32 males per session at T50, and 48 males at T67-69. At T50, 8 sessions (1 per day) were performed within 2 experimental blocks (*i*.*e*., independent age-cohorts created 6-7 days apart), for a total of 256 tested males. At T67-69, 15 sessions of observations were performed within 5 experimental blocks, for a total of 720 tested males.

### Post-mating pre-zygotic and early post-zygotic barriers: sperm precedence, fertilization failure and F1 hybrid mortality (Experiment 2)

To test for the evolution of post-mating pre-zygotic barriers (homotypic sperm precedence and fertilization failure) as well as early post-zygotic barriers (F1 hybrid mortality), a double-mating experiment was conducted following 56 to 64 experimental evolution transfers (‘T56-64’; see Box S2). Females were either mated with only one male (‘single mated females’) to assess fertilization failure and F1precedence, or with two different males (‘double-mated females’) to test for homotypic sperm precedence (*i*.*e*., preferential use of sperm from a male of their own form to avoid genetic incompatibilities; Zeh & Zeh, 1997). Only green-form females were used in this experiment (Figure 2B), because (1) fertilization failure was absent in crosses between red-form females and green-form males prior to experimental evolution (Cruz *et al*., 2025b), making deviation from first sperm precedence pattern more unlikely; (2) red-form females had undergone considerably fewer effective generations of selection than green-form females in the sympatry regime (see Box S2); and (3) red-form offspring are harder to distinguish from hybrids based on body coloration than green-form offspring (see Box S1). Testing red-form females would thus require the use of genetic markers to unambiguously assess paternity, an unaffordable technique given the high number of offspring produced.

Virgin females and males used in the experiment were obtained from age-cohorts created 11-12 days before the onset of the experiment. Briefly, *ca*. 50 green-form or 100 red-form mated females were transferred from the common garden of each replicate of each selection regime onto a fresh bean leaf placed on water-soaked cotton. Females were allowed to lay eggs for two days before being transferred to a new fresh leaf to lay eggs for another two days. Quiescent females and males were taken 5-6 days after the mothers were removed from each leaf discs, and kept separately on 4.9 cm^2^ leaf discs to ensure their virginity. Mating observations took place two days later, following eight experimental treatments: for each of the two selection regimes (sympatry or allopatry), green-form females were first mated with either a green- or a red-form male from the same regime (as above, from its own population replicate, or equivalent in the case of the allopatry regime). Then, half of the mated females was kept as ‘single-mated’ (females were not presented to any other male), whereas the other half was re-mated to a second male with the other colour form than the first male (females first mated with a green-form male could mate with a second red-form male, and females first mated with a red-form male could mate with a second green-form male; see Figure 2B).

For the first mating, 5 virgin females were placed with 5 males on 0.4 cm^2^ leaf discs (‘arenas’). Mating observations began as soon as the first male was installed with the 5 females and lasted for a maximum of 1 hour. Immediately after a mating occurred during this period, mated females were transferred to another arena, and males were discarded. Mated individuals were readily replaced by new virgin ones in the original patch. Females that did not mate at the end of the 1-hour observations period were discarded, and half of those that mated were transferred together to an empty 0.8 cm^2^ leaf disc. The remaining mated females were left on the arena, where new virgin males from the opposite colour form than the first males were added to launch the second mating. In this case, the procedure was the same as for the first mating except that the number of males installed per arena depended on the number of mated females obtained from the first mating to maintain a 50/50 sex-ratio. Then, mated females were immediately transferred to a 0.8 cm^2^ leaf disc after a mating occurred.

One day after the mating observations, single and double mated females were individually isolated on 2.5 cm^2^ leaf discs, where they laid eggs for 4 days and their daily survival was registered. On the fourth day, all females were discarded, and the eggs were counted. Unhatched eggs and juveniles were counted after four more days to obtain the egg hatching rate, and the number of dead juveniles, adult males, and green and hybrid daughters, were counted both six and eight days later. From patches with single mated females, we determined (a) fertilization failure potentially caused by inefficient sperm transfer/storage and/or gametic incompatibilities as the number of adult female *vs*. male offspring (offspring sex-ratio; as in Cruz *et al*., 2021), (b) F1 zygote mortality as the number of unhatched *vs*. hatched eggs, and (c) F1 juvenile mortality as the number of dead juveniles *vs*. adult offspring among hatched eggs. From patches with double-mated females, we tested for (d) homotypic sperm precedence by calculating the number of daughters sired by the first *vs*. the second male (determined as the number of green-form *vs*. hybrid females when the first male was green, or the reverse when the first male was red).

A total of 7 experimental blocks were performed, with alternating 7 or 21 days between two consecutive blocks such that the entire experiment lasted for *ca*. 5 months. Within a given block, four sessions of observations (one per population replicate) were performed in two consecutive days (hence two sessions per day), for a total of 28 sessions of observations across the whole experiment. All treatments (single-*vs*. double-mated females with red- or green-form males first, for each selection regimes) were performed simultaneously within each session. As not all females mated with the first or second male, the number of (single or double) mated females obtained per treatment per population replicate (N=4 for this experiments) varied between 0 and 7 within each block, and between 17 and 33 for the entire experiment, for a grand total of 860 female replicates obtained at the end of the experiment.

### Late post-zygotic barriers: F1 hybrid fertility and F2 hybrid breakdown (Experiment 3)

To test for the evolution of late post-zygotic barriers, the fertility of viable F1 hybrid females and the viability of F2 males (classical test for F2 hybrid breakdown in spider mites; De Boer, 1982; Vala *et al*., 2000; Sugasawa *et al*., 2002) were assessed following 47 experimental evolution transfers (‘T47’; see Box S2 for effective generations of selection). Prior to the experiment, mass F0 crosses were performed within and between the green- and red-form populations to obtain F1 hybrid females with a red- or green-form mother (thus a green- or red-form father, respectively), along with ‘parental type’ F1 female controls, for both the allopatry and the sympatry regime (Figure 2C). To this aim, age-cohorts were first created by transferring ca. 50 mated females from the common garden of each replicate and selection regime onto a bean leaf placed on water-soaked cotton to lay eggs for three days before being discarded. Seven days later, 20 quiescent females and 20 adult males (or 40 of each for crosses between green-form females x red-form males, as these crosses result in a strong underproduction of hybrid females; Cruz *et al*., 2021), of the same or different colour form depending on the treatment, but, as in previous experiments, always from the same selection regime and corresponding population replicate, were installed together on 8 cm^2^ leaf discs. Individuals could mate and females laid eggs for four days before being killed. Seven days later, F1 deutonymph and/or quiescent daughters were transferred to 1.8 cm^2^ leaf discs where they remained for two days to complete their last moult and emerge as virgin adults.

To assess F1 hybrid fertility, the young F1 virgin females were isolated on 2.5 cm^2^ leaf discs to lay eggs and their daily survival was registered. Three days later, the number of eggs was counted to determine the proportion of ovipositing F1 females (F1 female fertility) and their daily fecundity. At this time, ‘parental type’ females were discarded but hybrid females were given two more days to lay eggs, and their eggs were counted again, to obtain a sufficient sample size (number of replicates and number of eggs per replicate) for subsequent fitness measurements on F2 offspring from these crosses (because of the very low fecundity of F1 hybrid females; see Cruz *et al*., 2021). Finally, F2 (male) offspring survival (hence F2 hybrid breakdown) was assessed by counting the number of unhatched eggs and dead and alive juveniles five to seven days after the females were discarded.

The entire experiment was performed in 2 experimental blocks (independent age-cohorts) onset one day apart. Within each block, 12 F1 females were tested simultaneously per treatment and population replicate (N=5 for this experiment), for a total of 960 F1 females in the entire experiment.

### Statistical analyses

All statistical analyses were carried out using the R statistical Software (v. 4.4.2; R Core Team, 2024). The general procedure used to analyse the data was similar for all experiments and is fully detailed in supplementary Table S1. A maximal model including all fixed effects and their interactions was built for each response variable. The type of model and error family used for each analysis was chosen based on error distribution, and when several models fitted the data, the best model was selected based on marginal Akaike information criterion (mAIC). Minimal models were then established by sequentially removing non-significant variables from the maximal models, and the statistical significance of the explanatory variables was established using chi-squared tests (Crawley, 2007) with the Anova function (car package; Fox & Weisberg, 2019). The significant chi-squared values reported are for the minimal model while the non-significant values correspond to those obtained before removing the variable from the model. In all experiments, whenever fixed effects (alone or in interactions) were found to be significant, pairwise comparisons between the factor levels of interest were performed using z-tests (with the functions ‘emmeans’, then ‘contrast’ or ‘pairs”, emmeans package; Lenth *et al*., 2021). Whenever a given set of data was used in more than one comparison, the p-values of the pairwise comparisons were corrected with the Holm-Bonferroni method to account for multiple testing (i.e., classical chi-square Wald test for testing the global hypothesis *H*_*0*_; Holm, 1979).

When testing for pre-mating isolation, mating propensity and mate preference, data were computed as binary response variables and analysed using generalized linear mixed effect models (glmmTMB, glmmTMB package; Brooks *et al*., 2017) with a binomial error distribution. Latency to copulation and copulation duration were analysed using a cox proportional hazard mixed-effect model (coxme, coxme package; Therneau, 2015), a non-parametric method to analyse time-to-event data (Crawley, 2007). The selection regime, the colour form of the male and of the chosen female (for latency to copulation and copulation duration), the transfer from which the tested individuals originated (T50 or T67-69), as well as all interactions between these factors were fit as fixed explanatory variables, whereas the day at which each observation session took place and the population replicate were fit as random explanatory variables. In the specific cases where the experimental evolution transfer had a significant effect within an interaction, pairwise comparisons were performed separately within each transfer.

As above, mating propensity data (binary response variable; mated or not) obtained in the second experiment were analysed using a glmmTMB with a binomial error distribution. All other proportion data, testing for early post-mating isolation (*i*.*e*., F1 zygote and juvenile mortality, F1 sex ratio, and proportion of daughters sired by the first mate of double mated females, to assess sperm precedence) were computed by binding together the number of successes and failures (*e*.*g*., the number of females *vs*. males for sex ratio) with the function cbind, thereby accounting for the total number of observations (Crawley, 2007). These data were subsequently analysed using a glmmTMB with a (zero-inflated) betabinomial error distribution to correct for overdispersed errors (testDispersion, DHARMa package; Hartig, 2016). The colour form of the mate (or order of mates’ colour form) and the selection regime (allopatry or sympatry) of the tested green-form female, as well as the interaction between these two factors, were fit as fixed explanatory variables, whereas the population replicate and the experimental block were fit as random explanatory variables. For the analyses of F1 zygote mortality and F1 juvenile mortality, we built an additional model in which sex ratio (here directly computed as a proportion bounded between 0 and 1; *i*.*e*., number of daughters/number of adults) was added to the previous model as a fixed explanatory variable. By doing so, we tested whether using sex ratio as a covariate could improve the fit of the model and change the significance of the other explanatory variables.

Data testing for late post-zygotic isolation (*i*.*e*., hybrid fertility and hybrid breakdown), were also analysed with glmmTMB models. The proportion of F1 ovipositing females (i.e., females that laid at least one egg) was computed as a binary response variable and analysed using a binomial error distribution, whereas the total number of eggs laid by these ovipositing females was analysed using a Conway-Maxwell Poisson distribution (‘compois’ family in glmmTMB), which is parameterized with the exact mean (V = μφ), to correct for overdispersed errors (Huang, 2017). The hatching rate of F2 eggs was computed using the function cbind (number of hatched eggs versus number of unhatched eggs) and analysed using a binomial error distribution (or betabinomial in the case of overdispersion). The proportion of F1 ovipositing females and the number of eggs laid by fertile females were analysed for data obtained after 3 days of oviposition for all types of F1 females tested (i.e., resulting from all types of female x male F0 crosses: green x green, red x red, green x red and red x green), then also for data obtained after 3 and 5 days of oviposition for hybrid females (Holm-Bonferroni corrections were performed to account for multiple testing). The type of F0 cross, the selection regime, and the number of days of oviposition (in the analyses of hybrids only), as well as all interactions between these factors, were fit as fixed explanatory variables, whereas the population replicate, the experimental block, and the identity of each focal female (in the analyses of hybrids only due to repeated measures at day 3 and day 5) were fit as random explanatory variables. The hatching rate of eggs laid by ‘parental type’ and by hybrid F1 females were analysed separately, as those eggs were laid for different oviposition duration (3 or 5 days respectively). In both cases, the type of F0 cross and the selection regime, as well as their interaction, were fit as fixed explanatory variables, whereas the population replicate and the experimental block were fit as random explanatory variables (Table S1).

### Strength and contribution of each reproductive barrier to total isolation

The strength of reproductive barriers (*RI*_*n*_) across selection regimes can only be compared if these barriers are measured on a similar scale. Even though different methods have been employed for such an endeavour (*e*.*g*., Coyne & Orr, 1989; Mendelson, 2003; Ramsey *et al*., 2003), here we used two types of indices to maintain consistency with previous work on the same system (Cruz *et al*., 2021, 2025b).

For premating isolation (*RI*_1_), homotypic sperm precedence (*RI*_2_), and hybrid sterility (*RI*_5_), we used the same formulation as in Martin & Mendelson (2016):

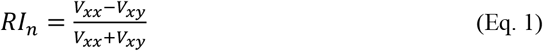

in which *V*_*xx*_ are the values observed in control homotypic crosses (i.e., from the allopatry regime only), and *V*_*xy*_ are those obtained from heterotypic crosses of a given regime, for each studied variable (*i*.*e*., number of copulations, offspring proportion in the brood of double mated females, number of eggs laid, for *RI*_1_, *RI*_2_, and *RI*_3_, respectively).

For fertilization failure (*RI*_3_), hybrid inviability (*RI*_4_), and hybrid breakdown (*RI*_6_), we used similar indices as those used in Cruz *et al*., (2021), adapted from Poinsot et al (1998). These indices compute the proportion of offspring affected by a given type of incompatibility (i.e., overproduction of sons and embryonic death of daughters in the case of fertilization failure and hybrid inviability in haplodiploids, and mortality of F2 zygotes in the case of hybrid breakdown) relative to the control crosses, thereby accounting for background variations:

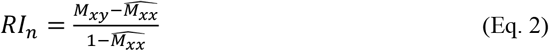

in which *M*_*xy*_ are the values observed in heterotypic crosses, and 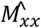 are the mean values observed in control homotypic crosses (i.e., from the allopatry regime).

These formulae, which are detailed for each barrier in Box S3, result in indices of isolation ranging from -1 to +1 (Eq. 1) or 0 to 1 (Eq. 2). A value of 0 indicates no isolation (i.e., no restriction in gene flow) due to the reproductive barrier, whereas positive and negative values indicate decreasing or increasing gene flow between species, respectively. Hence, here we considered that fertilization failure, hybrid inviability, and hybrid breakdown cannot be lower in control homotypic crosses than in heterotypic crosses, whereas negative values for premating isolation indicate disassortative mating (as observed between green females and red males in Cruz *et al*., 2025b), negative values for homotypic sperm precedence indicate that heterotypic sperm outcompetes homotypic sperm, and negative values for hybrid sterility would indicate heterosis (i.e., better fitness of hybrid than ‘parental type’ females).

Finally, we employed a method previously adapted from Coyne & Orr (1989) by Ramsey *et al*., (2003), in which total (cumulative) reproductive isolation between two populations or species is computed as a multiplicative function of the strength of each reproductive barrier (*RI*_!_), so that the contribution (*C*_*n*_) of each barrier to reducing gene flow at a stage *n* in life history is calculated as:

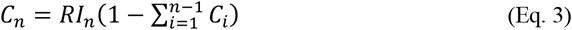

Thus, a given reproductive barrier eliminates gene flow that has not been prevented by earlier barriers. For example, fertilization failure (*C*_3_) acts later than premating isolation (*C*_1_) and could thus restrict a smaller share of gene flow than suggested by raw reproductive isolation strength (*RI*_3_). Here, we used the average of the *RI*_*n*_ values that were computed separately for each population replicate. Negative *RI*_!_ averages were set to 0, such that the contribution of all barriers to total reproductive isolation ranges from 0 to +1, indicating either a reduction in gene flow between populations (positive values) or equal amounts of gene flow within and between populations (zero values). Then, for *m* reproductive barriers, total reproductive isolation (*T*) between populations after experimental evolution was computed as:

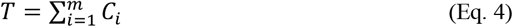

## Results

### Pre-mating barriers and mating behaviours (Experiment 1)

#### Mating propensity and mating preference

Overall, *ca*. 90% of the tested males mated during the 30 minutes of observation, independently of the tested generation (*X*^*2*^_*1*_*=2*.*51, p=0*.*11;* Model 1.1 in Table S1). However, only green-form males showed different mating propensity between selection regimes, with *ca*. 5% more males mating when they originated from the sympatry regime as compared to the allopatry regime (*male form*regime*: *X*^*2*^_*1*_*=3*.*75, p=0*.*05*; Figure 3A; see pairwise comparisons in Table S2). Yet, mating preference did not differ significantly between the two types of males in either of the selection regimes (*i*.*e*., the male form, as well as all its interactions with the regime or any other factor, were non-significant; Model 1.2 in Table S1). Most males, irrespective of their colour form, chose red-over green-form females (74% on average; Figure 3B, see also Table S3 for each replicate population). This preference, however, changed depending on the selection regime across generations (*generation time*regime*: *X*^*2*^_*1*_*=4*.*18, p=0*.*04*), as the proportion of (both red- and green-form) males that choose a red-form female decreased by more than 10% between the generation 50 and 67-69 in the sympatry regime, whereas it remained constant in the allopatry regime (see Table S2 for the comparisons).

**Figure 3.**
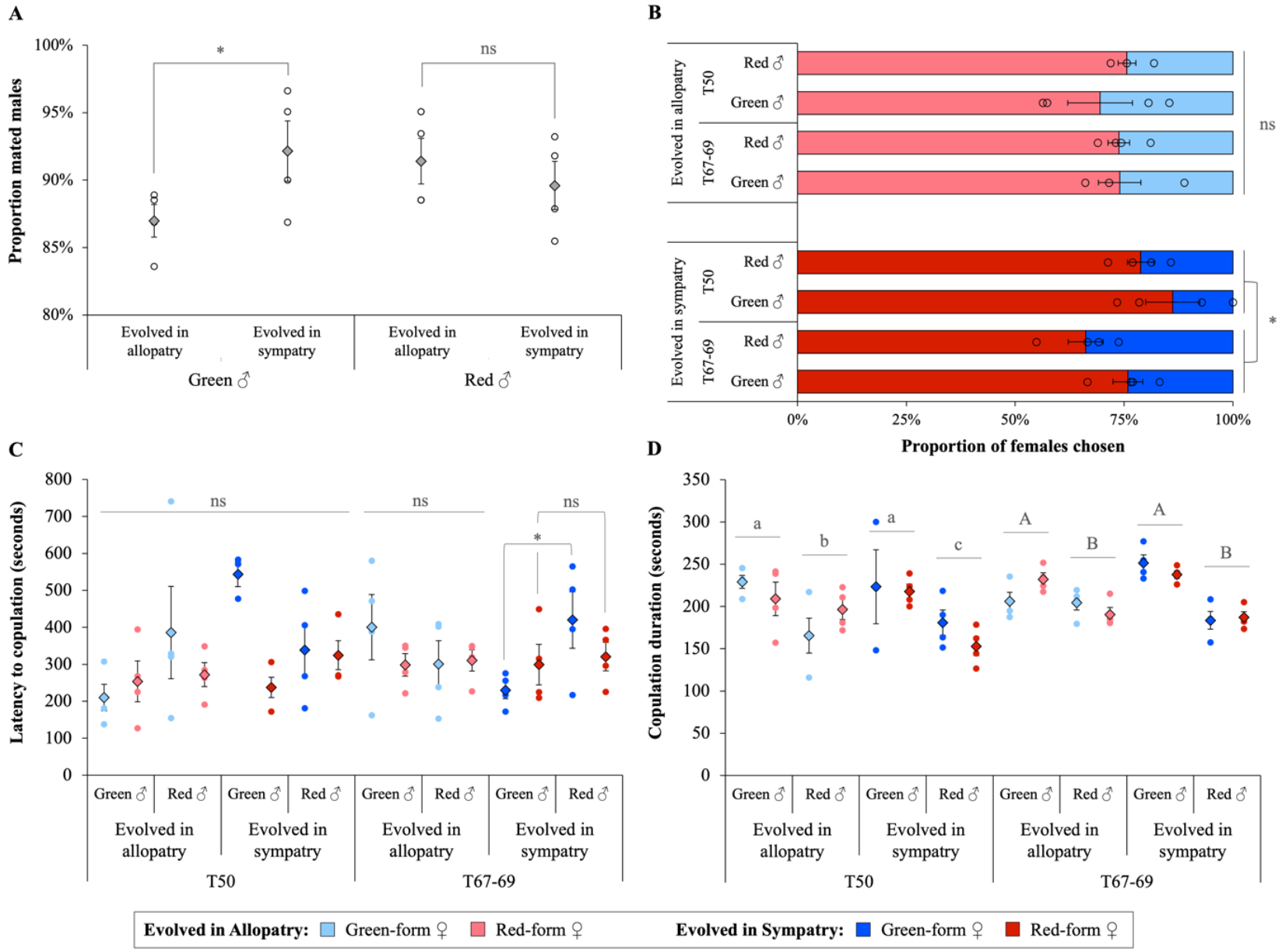
Mating behaviour of red- and green-form males after evolution in sympatry or allopatry (Experiment 1). (A) Mating propensity of males irrespective of the female they mated with, (B) proportion of females of each colour form chosen by each type of male, (C) latency to copulation, and (D) copulation duration (in seconds) of each type of male after 50 and 67-69 transfers of experimental evolution (“T50” and “T67-69”, respectively). Dots show the average trait value for each independent population replicate and diamonds (or bars in panel B) the grand means ± standard errors of these replicates. Boxplots showing the data distribution per replicate for latency to copulation and copulation duration are provided in Fig. S1. Lighter and darker colour tones represent individuals (both females and males) evolved in allopatry or sympatry, respectively. Blue: green-form females; Red: red-form females. ‘ns’ or identical superscripts indicate non-significant differences at the 5% level; *p<0.05.

#### Copulation latency and duration

Latencies to copulation were overall highly variable, ranging from ca. 20 seconds to almost 30 minutes across the entire experiment, and statistical analyses revealed a significant four-way interaction between the colour form of the males and of the chosen females, their selection regime and the generation at which they were tested (*X*^*2*^_*1*_*=5*.*61, p=0*.*02*; Model 1.3 in Table S1). Pairwise comparisons (Table S2) revealed that this complex interaction was due to green males mating *ca*. 3 minutes sooner with green females than red males, when they evolved in the sympatry regime for 67-69 generations (or transfers; Figure 3C; see also Figure S1 for each replicate population). Conversely, the duration of copulations did not differ between matings with green- and red-form females (*X*^*2*^_*1*_*=0*.*0007, p=0*.*98*) but varied between males of different colour forms depending on their selection regime, and across generations (*male form*regime: X*^*2*^_*1*_*=6*.*93, p=0*.*008; generation time*regime: X*^*2*^_*1*_*=4*.*37, p=0*.*04*; Model 1.4 in Table S1). Indeed, although green-form males consistently mated longer than red-form ones across selection regimes and generations (see Table S2 for comparisons between data obtained for males of different colour forms evolved in each selection regime for 50 or 67-69 generations), pairwise comparisons revealed that red males evolved in the sympatry regime mated for a significantly shorter time than those evolved in the allopatry regime, but only following 50 generations of experimental evolution, as this effect was lost after 67-69 generations (Figure 3D, and Table S2 for pairwise comparisons).

### Early post-mating barriers: F1 offspring mortality, fertilization failure and sperm precedence (Experiment 2)

#### Mortality of F1 offspring of single-mated green-form females

The mortality of F1 eggs (*i*.*e*., the proportion of unhatched eggs indicating *F1 zygote mortality*) was overall low (Figure 4A), and it was differently affected by the colour form of the male with which females had mated, depending on the selection regime (*male form*regime*: *X*^*2*^_*1*_=4.13, p=0.04; Model 2.2.1 in Table S1). Indeed, F1 zygote mortality was 2.6 ± 0.4% lower if females that evolved in sympatry had mated with a green-rather than a red-form male, whereas it remained unchanged for females evolved in the allopatry regime (see comparisons in Table S4). The opposite pattern was found for F1 juvenile mortality, with a higher increase in juvenile mortality in the brood of females that had mated with a green-rather than a red-form male from the sympatry regime (*male form*regime*: *X*^*2*^_*1*_=5.37, p=0.02; Model 2.3.1 in Table S1; Figure 4B). This led to a higher juvenile mortality of offspring from green females mated with sympatric vs allopatric green males (homotypic crosses), whereas no significant difference was found between the offspring of green females mated with red males (hence producing hybrid daughters) from the two selection regimes (see comparisons in Table S4). Moreover, further analyses revealed that adding sex ratio as a covariate in the two models significantly improved their fit and vanished the significant effects of the mate and selection regime of the tested females (see Models 2.2.1 *vs*. 2.2.2 and Models 2.3.1 *vs*. 2.3.2 in Table S1). Sex ratio explained most of the observed variance, as the proportion of females in the brood negatively correlated with zygote mortality (*X*^*2*^_*1*_=12.10, p<0.001; Model 2.2.2), but positively correlated with juvenile mortality (*X*^*2*^_*1*_=57.95, p<0.001; Model 2.3.2).

**Figure 4.**
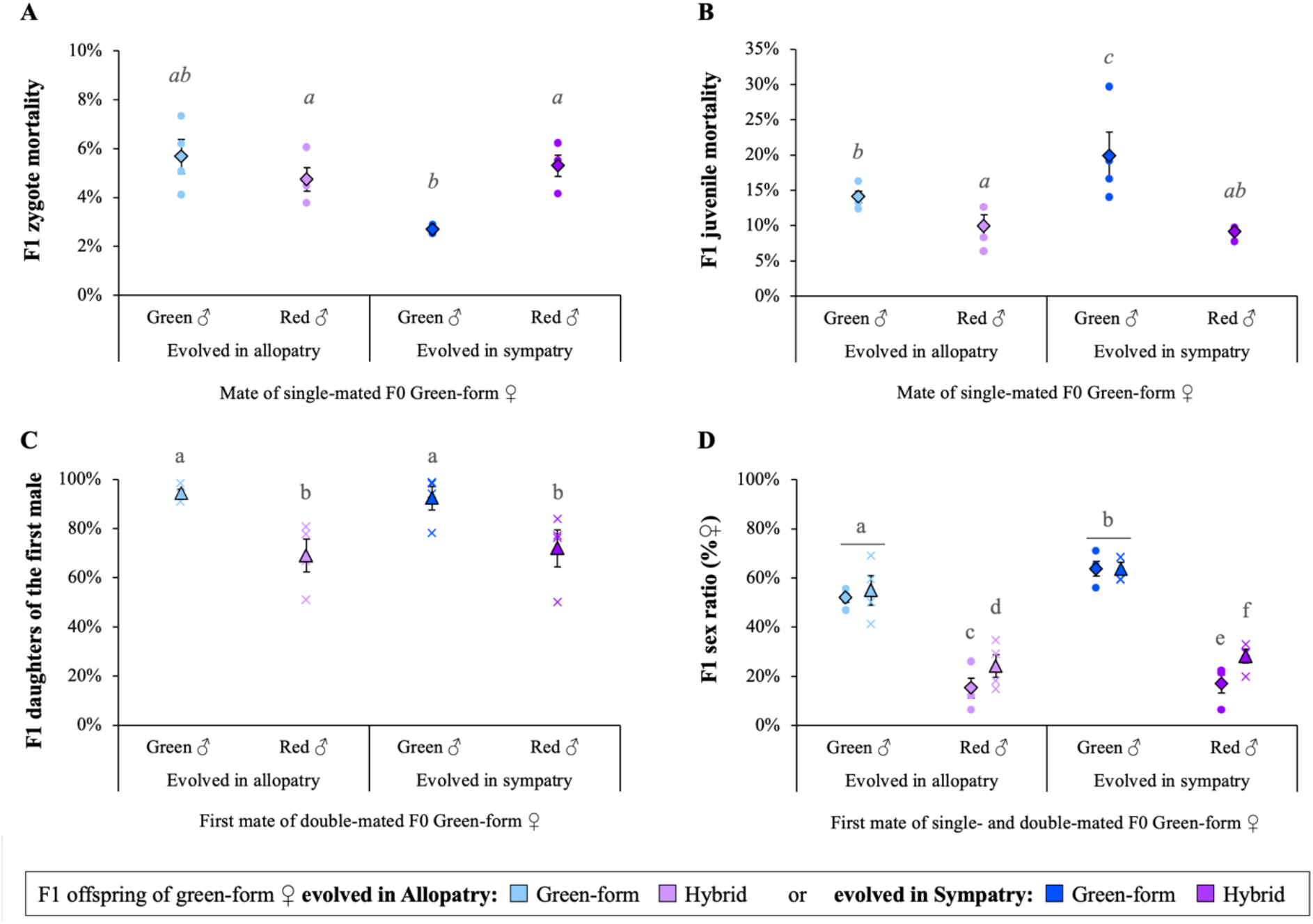
Fertilization failure, sperm precedence and F1 hybrid mortality after evolution of green-form females in artificial sympatry or allopatry (Experiment 2). (A) F1 zygote mortality (proportion of unhatched eggs) and (B) F1 juvenile mortality (proportion of hatched eggs that did not reach adulthood) in the brood of green-form females single-mated with a homotypic or heterotypic mates, (C) proportion of F1 daughters of the first mate (*vs*. those sired by the second mate) among the F1 female offspring of double mated F0 females, and (D) sex-ratio (proportion of daughters among adult F1 offspring) in the brood of single or double mated green-form females after experimental evolution. In all panels, the x axis indicates the selection regime and the colour form of the only (in A and B) or of the first (in C and D) male that mated with the focal female (in C and D second males were always from the other form than the first male). Dots and crosses show the average trait value for each independent population replicate or for single or double mated females, respectively. Diamonds or triangles show the grand means ± standard errors of all replicates for single or double mated females, respectively. Boxplots showing the data distribution per replicate are provided in Figure S2. Lighter and darker colour tones represent individuals (both females and males) from the allopatry and sympatry regime, respectively. Blue: homotypic crosses; Purple: heterotypic crosses. Identical superscripts indicate non-significant differences at the 5% level. In (A) and (B), superscripts are displayed in italics to indicate that significant differences depend on the statistical models used: no differences were found between types of mates when the sex ratio was accounted for as a covariate in the models.

#### Sperm precedence in double-mated green-form females

On average, 23 ± 5% fewer F1 females were sired by the first mate of green-form females when this mate was a heterotypic red-form male rather than a homotypic green-form male (ca. 70 ± 5% of F1 females sired by heterotypic first males *versus* 93 ± 2% sired by homotypic first males; *X*^*2*^_*1*_*=34*.*38, p<0*.*0001*; Model 2.4 in Table S1; Figure 4C). This shows that some degree of homotypic sperm precedence occurred despite the predominance of first male sperm precedence in this system. However, the amplitude of this effect did not vary between selection regimes (*male form*regime*: *X*^*2*^_*1*_*=2*.*27, p=0*.*13*; *regime effect*: *X*^*2*^_*1*_*=0*.*001, p=0*.*97*; Model 2.4).

#### Sex ratio and fertilization failure in single-mated versus double-mated green-form females

Whereas the F1 offspring sex ratio of green females mated first (or only) with a green male was slightly female-biased (58 ± 3% daughters among adult offspring on average), we found a drastic reduction in the proportion of daughters in the brood of green females first (or only) mated with a red male (21 ± 3% daughters among adult offspring on average; Figure 4D). These results, which corroborate those obtained previously with the same source populations (Cruz *et al*., 2021), generally indicate *fertilization failure* in spider mites, as daughters and sons are produced from fertilized and unfertilized eggs, respectively. In line with the pattern of sperm precedence described above, we found that this reduction in the proportion of daughters was weaker for females that remated with a homotypic (green) male after mating with a heterotypic (red) male, but not for females that remated with a heterotypic (red) male after a homotypic (green) one (*female status * colour form of the first/only mate*: *X*^*2*^_*1*_*=6*.*38, p=0*.*01*; Model 2.5 in Table S1; see comparisons in Table S4). Finally, the proportion of F1 females also differed between selection regimes (*effect of regime: X*^*2*^_*1*_*=5*.*84, p=0*.*02*; see Model 2.5 in Table S1), with more daughters being produced by females that evolved in sympatry than in allopatry (64 ± 2% *vs*. 53 ± 3% green F1 females, and 22 ± 3% *vs*. 19 ± 4% hybrid F1 females, respectively), but the amplitude of sex-ratio distortion (*i*.*e*., fertilization failure) observed in heterotypic *vs*. homotypic crosses, and in double *vs*. single mated females, was similar in both selection regimes (*colour form of the first/only mate*regime*: *X*^*2*^_*1*_*=2*.*00, p=0*.*15; female status*regime*: *X*^*2*^_*1*_*=0*.*33, p=0*.*57*; Model 2.5 in Table S1). Hence, although evolution in sympatry resulted in an increased production of daughters in heterotypic crosses (*i*.*e*., decrease in early post-zygotic barrier), the same increase in daughter production was also observed for homotypic crosses.

### Late post-zygotic barriers: F1 female fertility and F2 offspring mortality (hybrid breakdown) (Experiment 3)

#### Proportion of fertile F1 female and daily oviposition

The proportion of F1 ovipositing females differed depending on their parental cross (*effect of F0 cross*: *X*^*2*^_*3*_*=312*.*17, p<0*.*0001*), but it was not affected by the selection regime (*effect of regime*: *X*^*2*^_*1*_*=0*.*94, p=0*.*33*; *F0 cross*regime*: *X*^*2*^_*3*_*=1*.*79, p=0*.*62*; Model 3.1.1 in Table S1; Figure 5A). Indeed, the proportion of ovipositing females only differed between ‘parental type’ and hybrid F1 females (97 ± 1% of green or red-form females, against 9 ± 1% of the hybrids, laid at least one egg within 3 days), but no differences were found within these two groups (*i*.*e*., between green- and red-form females, or between hybrid females from different parental crosses; see all averages per replicate in Table S5 and comparisons in Table S6). Next, the number of eggs laid by fertile females over 3 days also varied depending on the type of F1 female (*effect of F0 cross*: *X*^*2*^_*3*_*=249*.*33, p<0*.*0001*; Model 3.2.1 in Table S1; Figure 5B), with similar number of eggs laid by green-form and red-form F1 females, but fewer eggs laid by hybrid females, especially those resulting from crosses between a green-form mother and a red-form father (see comparisons in Table S6). The number of eggs laid by all females, however, was overall lower when they originated from the sympatry regime as compared to the allopatry regime (*effect of regime*: *X*^*2*^_*1*_*=17*.*03, p<0*.*0001*), with no significant difference in the amplitude of this effect for hybrid and ‘parental type’ females (*F0 cross*regime*: *X*^*2*^_*3*_*=6*.*24, p=0*.*10*; Model 3.2.1 in Table S1; Figure 5C). Note also that, independently of the type of F0 cross and selection regime, the proportion of fertile F1 hybrid females did not change significantly if we considered 2 more days of oviposition (Model 3.1.2 in Table S1), although the number of eggs laid by the fertile hybrids increased significantly between 3 and 5 days of oviposition in the sympatry regime only (*regime*day*: *X*^*2*^_*1*_*=10*.*89, p<0*.*001*; Model 3.2.2 in Table S1; Table S5 and S6).

**Figure 5.**
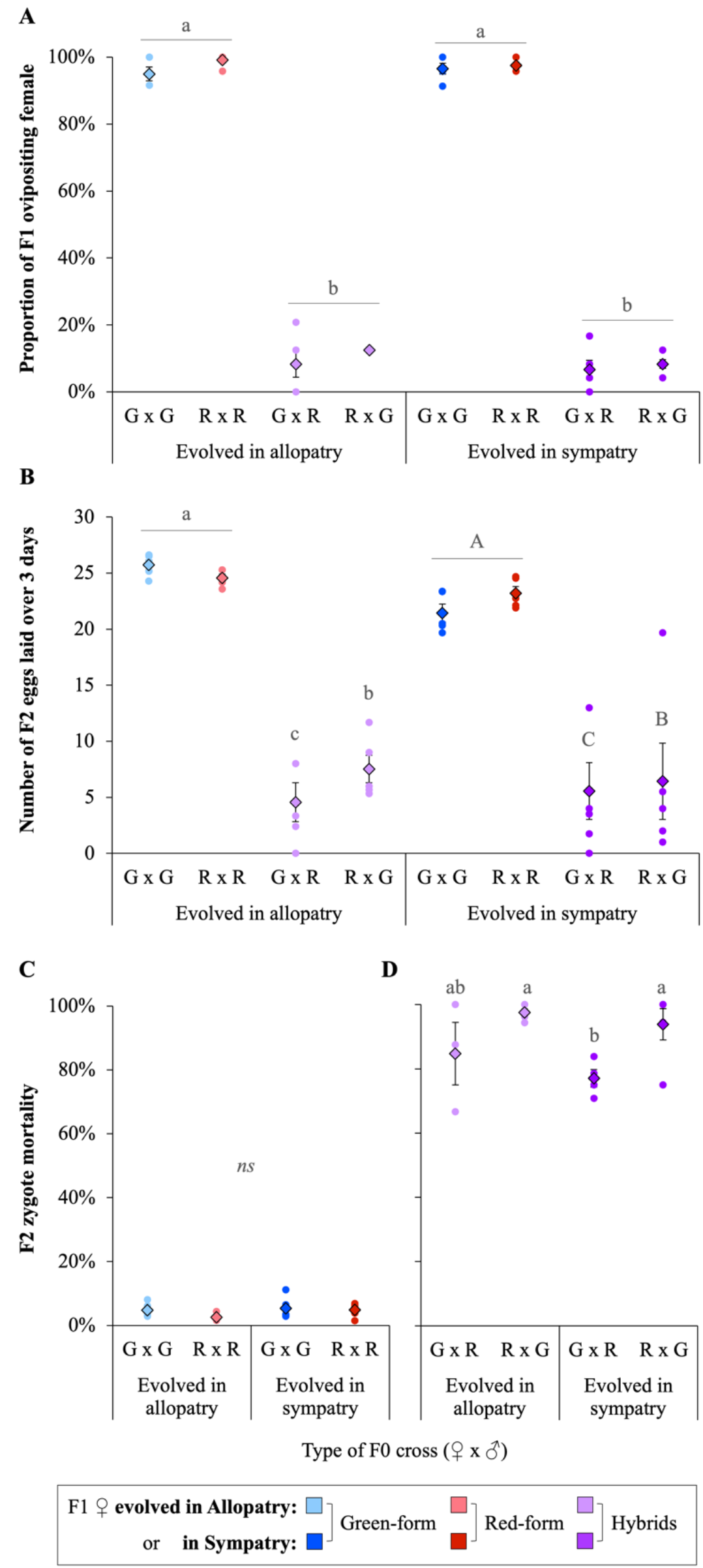
F1 hybrid fertility and F2 hybrid mortality after evolution in artificial sympatry or allopatry (Experiment 3). (A) Proportion of ovipositing F1 virgin females, (B) number of eggs laid over 3 days by these ovipositing females (i.e., excluding fully sterile females), (C) F2 zygote mortality (proportion of unhatched eggs) in the brood of F1 green- or red-form females, and (D) in the brood of F1 hybrid females, for different types of F0 crosses between green (G) and red (R) form F0 females and males (♀ x **♂**) after experimental evolution. Dots show the average trait value for each independent population replicate, whereas diamonds show the grand means ± standard errors of these replicates (boxplots showing the data distribution per replicate for F2 zygote mortality are provided in Figure S3). Lighter and darker colour tones represent crosses between individuals (both females and males) evolved in the allopatry or the sympatry regime, respectively. Blue: F1 green females (i.e., resulting from homotypic crosses between green F0 females and males); Red: F1 red females (i.e., resulting from homotypic crosses between red F0 females and males); Purple: F1 hybrid females (i.e., resulting from F0 heterotypic crosses). Identical superscripts or ‘*ns*’ indicate non-significant differences at the 5% level.

#### F2 zygote mortality

Whereas the proportion of unhatched F2 eggs in the brood of green and red-form females was very low (4.4 ± 0.7% on average) regardless of their colour form and selection regime (*F0 cross*regime*: *X*^*2*^_*1*_*=2*.*34, p=0*.*13*; *F0 cross*: *X*^*2*^_*1*_*=2*.*51, p=0*.*11*; *regime*: *X*^*2*^_*1*_*=0*.*11, p=0*.*74*; Model 3.3.1 in Table S1), it was much higher in the brood of hybrid females (90.2 ± 3.1% on average), and varied depending on the type of parental cross and selection regime from which these females originated (*F0 cross*regime*: *X*^*2*^_*1*_*=3*.*89, p=0*.*05*; Model 3.3.2 in Table S1; Figure 5D). Pairwise comparisons revealed that the mortality of F2 eggs laid by the two types of hybrid females differed when they originated from the sympatry regime, being ca. 16.8 ± 5.6% lower when their mother was a green female, but did not differ between the two types of hybrid females when they originated from the allopatry regime (see comparisons in Table S6).

### Strength and absolute contribution of each reproductive barrier to total isolation

Overall, we only found weak changes in the strength of all reproductive barriers in the sympatry regime relative to the allopatry regime, with no significant change in almost all reproductive barriers, including that involving changes in sex ratio (i.e., fertilization failure), except for a decrease in premating isolation in crosses between green-form females and red-form males (Figure 6A). As a result, only the contribution of this barrier to total reproductive isolation between green females and red males decreased in the sympatry regime, without any notable consequences for total reproductive isolation (99.88% *vs* 99.84% in the sympatry *vs*. allopatry regime; Figure 6B).

**Figure 6.**
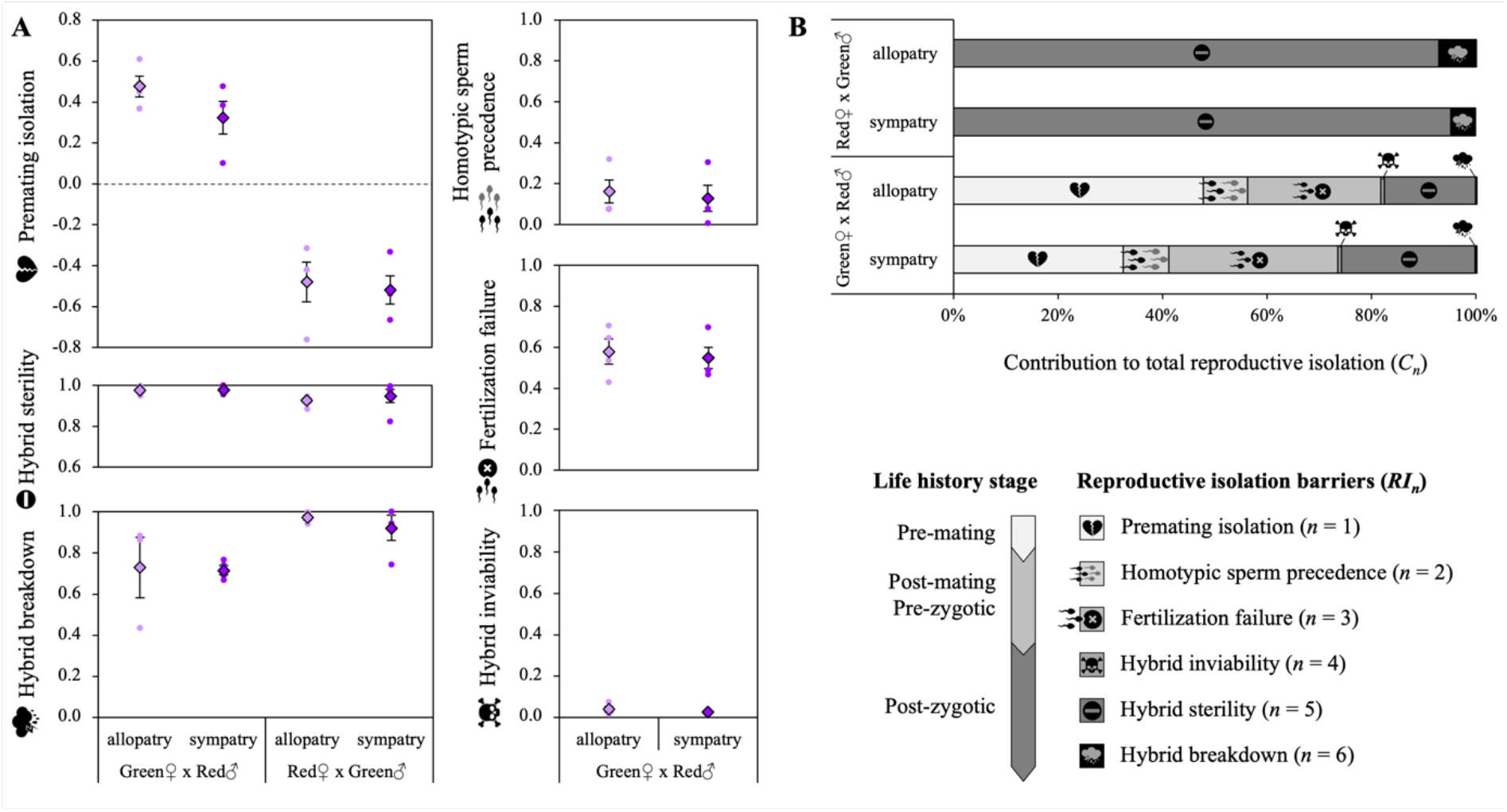
Estimation of (A) the strength of reproductive barriers and (B) their contribution to reducing gene flow between populations evolved in artificial sympatry or in allopatry. In (A), dots show the estimated strength of each reproductive isolation barrier (*RI*_*n*_) per population replicate, whereas diamonds show the average (± standard error) for each selection regime. Lighter and darker purple tones represent barriers occurring in heterotypic crosses between individuals (both females and males) evolved in the allopatry or the sympatry regime, respectively. In (B), stacked bars show the percent contribution of each barrier (*C*_*n*_) to total reproductive isolation (*T*). Three reproductive barriers (homotypic sperm precedence, fertilization failure and hybrid inviability) are not shown for crosses between red ♀ and green **♂**, as we did not assess their evolution (these barriers were absent prior to experimental evolution; Cruz et al. 2025).

## Discussion

Despite very high costs of hybridization and virtually inexistent gene flow between two closely related spider mite colour forms, long-term experimental evolution in artificial sympatry *vs*. allopatry did not lead to reinforcement, even though the relative strength of some reproductive barriers changed. Indeed, the preference of both green- and red-form *T. urticae* males for red-form females decreased when the two colour forms evolved in sympatry (leading to decreased disassortative and assortative mating, respectively), whereas it remained constant when they evolved in allopatry. This change was accompanied by an overall increase in the mating propensity of green-form males, and a decrease in their latency to copulation with green-form females.

Next, our investigation of offspring production by single- and double-mated green-form females showed that first male sperm precedence is incomplete in this system, especially when these females first mated with a heterotypic male. This strongly suggested that the sex-ratio shift towards males occurring in the offspring of green-form females mated with red-form males (Cruz *et al*., 2021) is due to fertilisation failure (Potter & Wrensch, 1978; Satoh *et al*., 2001; Costa *et al*., 2023; Cruz *et al*., 2025b). Yet, the sex ratio in the offspring of such females already mated with a heterotypic male only becomes slightly less male biased when they subsequently mated with a homotypic male (see also Cruz *et al*., 2025b). This work further revealed that green-form females originating from the sympatry regime produced more hybrid daughters than those from the allopatry regime (both in single matings with red males and after a second mating with a green male), suggesting a decrease in post-mating pre-zygotic barriers in sympatry. However, a similar increase in daughter production was found when these females engaged in homotypic crosses, thus revealing that sex allocation evolved in sympatry irrespective of the type of male these green-form females mated with. Moreover, this outcome had cascading consequences on F1 zygote and juvenile mortality as both positive and negative correlations were found between these vital rates and offspring sex ratio.

Then, we confirmed that late post-zygotic barriers are nearly complete in this system, as crosses in both directions generated hybrids with very low fitness (Cruz *et al*., 2021). Yet, evolution in sympatry had little impact on those patterns, with only slight fitness changes in F1 virgin females (*i*.*e*., lower fecundity but higher F2 embryonic mortality), irrespective of whether they were hybrids or not.

Finally, compiling these results confirmed the strong asymmetries in reproductive barriers between these forms (as in Cruz *et al*., 2025b), and shows that, running counter predictions of reinforcement (Gavrilets, 2014), most barriers did not substantially change following evolution in artificial sympatry, and some of them even tend to decrease, *i*.*e*., mate preference and fertilization failure in crosses between red-form males and green-form females.

Given that reproductive interactions between the green and red colour forms studied here are highly costly, we hypothesized that evolving in sympatry would either lead to stronger pre-mating isolation or to fewer post-zygotic incompatibilities. Instead, responses to the selection regimes were in general weak. Several hypotheses can be put forward to explain this. First, even though previous experimental evolution with spider mites showed that evolutionary responses can occur within few generations (*e*.*g*., Magalhães *et al*., 2007a; Belliure *et al*., 2010; Bitume *et al*., 2011; Fragata *et al*., 2025), it may be that these population have not evolved together for sufficient time (Fig. II in Box S2). Second, while selection might be stronger in the red-form population because red-form females are preferred over green-form ones (hence should suffer the most), the red-form population is the inferior competitor in our lab conditions (Cruz *et al*., 2025a). Although we compensated for this by introducing more red-form individuals in the sympatric populations, reduced population size and limited genetic variation in response to competition in artificial sympatry may thus have limited their evolutionary potential (Johansson, 2008; Osmond & De Mazancourt, 2013; Chen & Zhang, 2024). However, none of these explanations is compatible with the fact that we observed evolutionary responses, sometimes in an opposite direction to that expected due to reinforcement. Instead, traits convergence and evolution of decreased reproductive isolation could potentially result from gene flow occurring between the studied populations (Grant *et al*., 2004; Grabenstein & Taylor, 2018; Birzu *et al*., 2025), especially given that we found lower hybrid sterility here than in previous work (more than 99% of hybrid females were found fully sterile in Cruz *et al*., 2021 *vs. ca*. 90% here). However, in both cross directions, (the few) hybrid females that were fertile consistently had very low fecundity and their eggs showed very low hatchability. Given such high costs of hybridization, we believe that gene flow is not a likely explanation for the evolution of decreased reproductive barriers in this system.

Whereas most changes in reproductive barriers, as well as the other responses observed after evolution in sympatry in our study, cannot be explained by selection against hybridization, they may still be indirect within-species responses to the consequences of sympatry. In particular, it has been argued that within-species, sexual selection and sexual conflict may indirectly affect the evolution of reproductive barriers between species in sympatry (Ritchie, 2007; Maan & Seehausen, 2011; Gavrilets, 2014). Although these forces are traditionally studied for their role in driving the evolution of reproductive isolation, both theoretical and empirical studies suggested that they can also constrain population divergence and inhibit speciation (*e*.*g*., Magurran, 1998; Parker & Partridge, 1998; Gavrilets & Waxman, 2002; Kirkpatrick & Nuismer, 2004; reviewed in Ritchie, 2007; Svensson *et al*., 2009), sometimes even driving the evolution of stronger reproductive interference (i.e., costly sexual interactions between species; Kyogoku & Sota, 2015, 2017; Yassin & David, 2016).

In our study, variable degrees of sexual conflict over mating rate initially present in the studied populations could explain the patterns of assortative and disassortative mating observed in allopatry, as well as their decrease after evolution in sympatry. In allopatric conditions, one could expect sexual conflict over mating rate to be stronger in the green-form population than in the red-form population because the sex-ratio of the green-form population is more male-biased at high population density (see supplementary Box S4), and sexual conflicts generally increase with the frequency of males within populations (McDonald *et al*., 2025; but see Chmielewski *et al*., 2026). Moreover, the fact that green-form males consistently mate longer than red-form males (here in all tested scenarios, and throughout three independent experiments in Cruz *et al*., 2025b) suggests that male-male competition is more intense in the green-form population. With an increase in the intensity of sexual conflict over mating rate, females are expected to evolve strategies to avoid coercive males (Smit, 2025). In particular, being less attractive could allow females to avoid excessive matings, as sexual attractiveness can be costly in spider mites (Li & Zhang, 2021). This could explain the reduced attractiveness of green-form females, hence why red-form ones are preferred by both types of males (Cruz *et al*., 2025b). In sympatric conditions, however, green-form females may benefit from relaxed sexual harassment due to the presence of red-form females (see Brask *et al*., 2012). A similar outcome has been found in two closely-related damselfly species (*Calopteryx splendens* and *C. virgo*) where competition between con- and heterospecific males benefited females by reducing mating harassment from conspecific males (Gomez-Llano *et al*., 2018). In our study, relaxed sexual harassment on green-form females in sympatry might thus have relaxed the pressure for maintaining low attractiveness, whereas attractiveness became very costly in red-form females. This may thus potentially explain the shift toward less preference for red-form females in both male types after evolution in sympatry.

Another main outcome of heterotypic interactions in sympatry is the overproduction of green males resulting from crosses between green-form females and red-form males. Such a reproductive interference-driven male-biased sex ratio should lead to increased male-male competition in sympatry as compared to allopatry, with several potential consequences. First, this may explain the evolution of increased mating propensity in green-form males (as observed in other species; Chechi *et al*., 2022), as well as the evolution of a reduced mating latency when these males mate with green-form females, allowing them to match the trait values of their red-form male competitors. In line with this, previous studies highlighted that evolution in sympatry can lead to phenotypic convergence instead of divergence, even in the absence of gene flow (Muschick *et al*., 2012; Weber & Strauss, 2016). Second, the increased proportion of males in the offspring may lead to selection for the production of more daughters, as they will have a higher reproductive value than sons. In line with this, recent theoretical work shows that, in haplodiploids, the presence of females constrained to produce only sons in a population (*i*.*e*., unfertilized females), favours the production of more female-biased offspring sex-ratios by unconstrained females (Godfray, 1990; Chokechaipaisarn & Gardner, 2025). Even though green-form females are not fully constrained in sympatry in our study (*i*.*e*., they still produce a few sterile hybrid daughters), the over-production of males in heterotypic crosses may have driven the evolution of sex allocation toward more female-biased offspring sex ratios. Moreover, the fact that this same outcome was found irrespectively of the type of males the green-form females mated with, further suggests that sex allocation is under female control, which is congruent with earlier studies in the spider mite system (Macke *et al*., 2011a, 2012a; Vala *et al*., 2003, but see Macke *et al*., 2014). Importantly, this also shows that the observed changes in sex allocation are not due to traits affecting fertilization success specifically in heterotypic crosses (e.g., traits involved in sperm transfer, storage or in sperm-egg compatibility), nor due to changes in traits affecting sperm competitive ability and/or cryptic female choice, as no changes in sperm precedence pattern were found when using mates evolved under sympatric conditions. This further suggests that traits involved in true changes in reproductive barriers may be under strong stabilizing selection, as they likely also directly affect homotypic crosses (Servedio & Hermisson, 2020).

Finally, several evolutionary responses observed after experimental evolution in sympatry could be easily explained by trait correlations due to trade-offs in resource acquisition and allocation. For instance, our results show that the changes in juvenile mortality in sympatry directly resulted from a positive correlation between this vital rate and the proportion of daughters in the brood of green-form females. This may be simply due to the fact that developing females may suffer more intense competition for food than males (which are much smaller) in this species (see Cruz *et al*., 2025a and Bonte & Bafort, 2019; Fragata *et al*., 2022; Fig. IE in Box S1), hence the higher the proportion of females, the higher the juvenile mortality. Inversely, we found a negative correlation between embryonic mortality (*i*.*e*., proportion of unhatched eggs) and the proportion of daughters in the brood of mated green female. This suggests better hatching of female embryos, probably because mothers allocate more resources to female eggs (as they are bigger than male eggs; Macke *et al*., 2011). Alternatively, both these patterns are also compatible with inbreeding depression in haplodiploids, as hemizygous males should suffer more from deleterious mutation; *i*.*e*., the positive correlation between adult female proportion and juvenile mortality may thus result from a higher mortality of male juveniles (but see Tien *et al*., 2015). Then, we also found that fertile RxG hybrids laid more eggs than fertile GxR hybrids, but their embryonic survival was worse (and a similar tendency can be observed for virgin green-form and red-form females; Figure 5). This may be indicative of asymmetrical Bateson-Dobzhansky–Muller incompatibilities, such as cytonuclear interactions, affecting different life stages (Barnard-Kubow *et al*., 2016; Knegt *et al*., 2017). However, the observed negative correlation between fecundity and hatch rate could also be due to a trade-off between the number of eggs laid by females and the quantity of resources allocated to each one of them. In line with this, a previous study showed a negative correlation between the size and the number of eggs laid by virgin females in spider mites (Macke *et al*., 2012b). Such a trade-off could also explain why virgin females originating from the sympatry regime overall laid fewer eggs than those originating from the allopatry regime, if evolving in sympatric conditions selected for producing bigger eggs, with a better embryonic survival and a higher likelihood of being fertilized (for mated females), hence to develop as a female. This work therefore unravelled several correlations among traits, which could underlie the evolutionary response to the presence of the other species.

## Conclusion

Our findings show no reinforcement between populations of the two closely-related colour forms of *T. urticae* under forced long-term evolution in sympatry. Instead, and despite very high costs of hybridization and very low (if not inexistent) gene flow, we observed an indirect (although weak) decrease in the relative strength of some reproductive barriers. However, trait changes observed in a given direction of heterotypic crosses were always accompanied by similar changes in reciprocal heterotypic crosses (*e*.*g*., for mate preference), or in the control homotypic crosses (*e*.*g*., for sex ratio). This suggests that the observed evolutionary responses, although driven by heterotypic interactions, mostly occurred at the intrapopulation level, with cascading effects on heterotypic reproductive interactions. More specifically, our results suggest that sexual conflict over mating rate and sex allocation play an important role in the maintenance of partial reproductive isolation (Burdfield-Steel & Shuker, 2011; Yassin & David, 2016; Kyogoku & Sota, 2017; Servedio & Hermisson, 2020), driving convergence rather than divergence between species traits in this system. In line with recent work, our findings show how the interplay between different types of interactions, within or between sexes of the same or different reproductively isolated populations (or incipient species), can shape evolutionary trajectories, and highlights the importance of considering higher degrees of complexity to understand unexpected evolutionary outcomes.

Finally, one notable prediction stemming from these results might be that only weak changes should be observed in the strength of reproductive interference (see Cruz *et al*., 2025a) following evolution in sympatric conditions. Support for such a prediction in the field could be stronger reproductive isolation between spider mite populations collected in allopatry than in sympatry (*e*.*g*., due to isolation by distance; see Sousa *et al*., 2019). However, other types of pre-mating barriers, which were prevented here, could also evolve in sympatry, such as divergence in habitat preference (incl. host plant specialization; Magalhães *et al*., 2007b; Sousa *et al*., 2019; Villacis-Perez *et al*., 2021), such that spider mites may speciate mostly via ecological speciation.

## Data Availability Statement

All datasets and R scripts used in this study are available at Zenodo (https://doi.org/10.5281/zenodo.18487013).

## Authors’ contributions

MAC, VS, SM and FZ conceived and designed the experimental evolution. MC and MAC performed the experimental evolution with the help of IS. MC, LRR and FZ conceived and designed the experiments to assess the evolution of the different reproductive barriers, and MC performed the experiments. FZ analysed the data and wrote the first version of the manuscript with input from MC and LRR. Subsequent versions also include inputs from MAC, VS and SM. LRR and FZ supervised MC. VS, SM and FZ supervised MAC. Funding agencies did not participate in the design or analysis of experiments. All authors read and approved this version of the manuscript.

## Acknowledgements

We are grateful to Cátia Eira and Lucie de Sousa for the maintenance of the spider mite populations and the plants, and to Bastien Aubry for his help in the mate preference experiment. We also thank all members of the Mite2 team at CE3C in Lisbon for useful advises with experimental designs, David Shuker and Alexandre Blanckaert for useful suggestions on earlier versions of the manuscript. This work was funded by an ERC Consolidator Grant COMPCON (“Competition under niche construction”, GA 725419) to SM, by the ERC Portugal Program of FCT, via the project DOFLOW (“Impact of Dominance and Gene flow on Adaptation”) granted to VS, the FCT project “GeneticBarriers” (2022.03475.PTDC) granted to Alexandre Blanckaert, and through the FCT strategic projects UIDB/00329/2020 granted to CE3C (https://doi.org/10.54499/UIDB/00329/2020). MAC was funded through an FCT PhD fellowship (SFRH/BD/136454/2018). FZ acknowledges support from CNRS and University of Montpellier. This is contribution ISEM-2026-XXX of the Institute of Evolutionary Science of Montpellier (ISEM). For the purpose of Open Access, a CC-BY 4.0 public copyright licence has been applied by the authors to the present document and will be applied to all subsequent versions up to an Author Accepted Manuscript arising from its submission. The authors declare no conflicts of interest.

## Notes

https://doi.org/10.5281/zenodo.18487013

### Competing Interest Statement

The authors have declared no competing interest.

### Summary of Updates

Analyses and results of mating propensity in Experiment 2 were removed from all files (Main file, Supplements, R Script and raw data), as these data could not be unambiguously attributed to mating propensity (i.e., missing data were erroneously interpreted as resulting from non-mated females).

