## supplementary for "Unexpected changes in reproductive barriers between incipient species after experimental evolution in sympatry"

### Box S1. Detailed information concerning the *T. urticae* source populations used in the study

All field and genetic information concerning the green- and red-form source populations used in this study, gathered throughout our previous studies [1-6], is summarized in Table I. These populations, called 'AMP' and 'TOM' in [1-4], 'Gu' and 'Ru1' in [5], then 'Gu' and 'Ru' in [6], were collected from locations *ca.* 34 km apart in central Portugal. They were established in the laboratory from 300 green females collected on tomato plants (*Solanum lycopersicum*) in May 2010, and 65 red form females collected on thorn apple plants (*Datura stramonium*) in November 2013, respectively. Following laboratory rearing, both populations were confirmed to be uninfected by any known endosymbiont of spider mites (*Cardinium*, *Rickettsia*, *Spiroplasma* and *Arsenophonus*; [2 ,4]), except *Wolbachia*, an endosymbiont capable of manipulating its host's reproduction [7]. Hence, to obtain *Wolbachia*-free green- and red-form populations, they were treated with rifampicin antibiotics for one generation in March and May 2018, respectively, as described in [4, 5]. Briefly, 400 adult females of each population were installed to lay eggs in petri dishes containing bean leaf fragments placed on cotton soaked in a rifampicin solution (0.05%, w/v), and from these eggs, 265 green and 308 red adult females were obtained to build the source populations used in this study. The absence of *Wolbachia* was confirmed three generations later though multiplex PCR diagnostic as described in [8].

**Table I. Information relative to the spider mite populations used in this study.** The table provides ITS2 and COI GenBank accession numbers matching the obtained sequences with 100% coverage and identity at the nucleotide level.

| Population form | Red | Green |
|---|---|---|
| **Original name** | AMP | TOM |
| **Collection date** | 18/11/2013 | --/05/2010 |
| **Host plant** | *Datura stramonium* | *Solanum lycopersicum* |
| **Location** | Aldeia da Mata Pequena | Carregado |
| **Coordinates** | 38.534363, -9.191163 | 39.078962, -8.993656 |
| **Number of sampled females** | 65 | 300 |
| **ITS2 GenBank No.** | GU565314 | AM408031 |
| **COI GenBank No.** | MF428440 | HM486513 |
| **References** | [1, 2] | [3, 4] |

DNA extractions from pools of 100 females for each population, and subsequent sequencing of a fragment of the nuclear ribosomal DNA *ITS2* (internal transcribed spacer 2) region and of the mitochondrial DNA Cytochrome Oxidase subunit I (*COI*), were performed as described in [1]. The results confirmed that both populations belong to *T. urticae*, but they also revealed that the two populations differ by 1 SNP in their *ITS2* sequence and by 21 SNPs in their COI sequence (genetic distance of 0.056 with Kimura 2-parameter; rate of variation gamma distribution; shape parameter = 1; [5]). This indicates a low level of genetic differentiation between the two populations based on these genetic markers (but see [9] for genomic differentiation between other populations of the two colour forms based on whole genome sequencing).

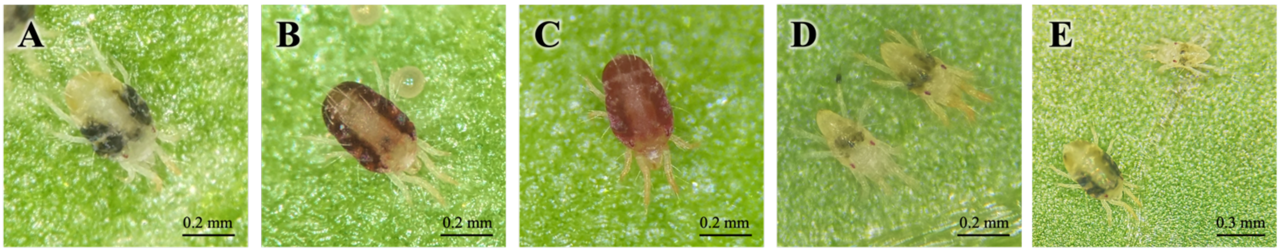

**Figure I. Typical body colours and sexual dimorphism in the *T. urticae* populations used in this study.** (A) green form, (B) red form, and (C) F1 hybrid females differ in their typical body colour, although hybrid females can sometimes be mistaken with red-form females. Conversely, the body colouration of (D) green-form (bottom left) and red-form (top right) males do not differ. (E) Adult *T. urticae* females (bottom left) and males (top right) can be easily distinguished owing to clear sexual dimorphism (incl. large body size difference). Figure also in [10].

Yet, previous studies showed that these two populations suffer strong but mostly asymmetrical reproductive isolation: (i) heterospecific crosses between green-form females and red-form males result in 75% overproduction of haploid sons instead of hybrid daughters, whereas the reciprocal cross does not lead to sex ratio distortion [5]; (ii) hybrid females (see Figure I) are fully sterile in both directions of crosses, with ca. 99% nonovipositing females and complete hybrid breakdown (none of the eggs hatched) [5]; (iii) females of the two colour forms do not show mate preference (random mating), whereas males of both forms prefer mating with red females (both assortative and disassortative mating) [6]; (iv) females mated with a heterotypic male (hence incompatible) are more attractive and/or receptive to subsequent homotypic males (hence compatible) than females that mated with a homotypic male. Yet, no clear benefit of this behaviour was found for offspring production (i.e., no homotypic sperm precedence) [6].

### Box S2. Experimental evolution procedure and effective generations of selection

#### *Modifications of the experimental procedure during the course of experimental evolution*

During the course of experimental evolution, the maintenance of the red-form populations in the sympatry regime was laborious, due to two main reasons: (1) the green-form populations were excluding the red-form populations through reproductive interference and/or resource competition [1], and (2) a proportion of the red females transferred at each generation might have been hybrids (*i.e.,* females produced from heterotypic crosses) instead of 'purebred' females (*i.e.,* females produced from homotypic crosses) due to their similarity in body colouration (Figure I in Box S1). To prevent the loss of replicates, missing red-form females were occasionally transferred from replicates of the allopatry regime (*e.g.,* females from replicate 1 of the red form allopatry regime were transferred into replicate 1 of the sympatry regime). However, this solution was problematic as it creates gene flow from populations of the allopatry regime into populations of the sympatry regime, thereby leading to non-complete independence of the evolution regimes and reducing the effective number of generations of selection in the sympatry regime (see next section below).

As a first attempt to avoid this issue, the experimental evolution procedure for the sympatry regime was modified in July 2019 (after 17 transfers): 400 red females (instead of 200) were transferred per replicate population, along with the usual 200 green females ('Procedure 2'; see Figure II). However, this solution was not sufficient, as red-form females were still rare in the sympatry regime and had to be taken from the allopatry regime at some occasions. As a result, one replicate was lost in December 2020 (after 53 transfers; Figure II).

We then implemented an additional modification of the experimental evolution procedure in January 2021 (after 55 transfers). This procedure ('Procedure 3'; Figure II), adapted from [2], consisted in the creation of a 'backup population' ($t - 1$) of red-form mites for each replicate of the sympatric regime at each transfer. Briefly, in addition to the regular transfers for each of the four remaining replicates of the sympatry regime, additional 200 to 400 red females were transferred to new boxes with two fresh plants without green females (*i.e.,* relaxed selection). If not enough red females were found in the sympatry replicate populations at the next transfer, the missing ones were taken from the corresponding $t - 1$ backup populations. In some cases, red females from the backup populations were maintained for up to four generations in absence of green females, such that females could be taken from these other backups when the sum of females found in the experimental population and its respective $t - 1$ backup did not add up to 400. This last modification of the procedure finally allowed successful maintenance of the sympatry regime without any further gene flow from the allopatry regime.

#### *Estimating the effective number of generations of selection*

Because, at every generation, each replicate population of the sympatry regime was replenished with variable numbers of migrants from the replicate populations of the allopatry regime, or from different backup populations, the number of effective generations of selection in this regime did not always correspond to the number of transfers performed, and differed for each replicate population. The effective number of generations of selection was estimated for each replicate population using the following equation, adapted from [2]:

$$Gen_{t+1} = S_t + \frac{N_t * Gen_t + \sum_{i=1}^{4}(N_{t-i} * Gen_{t-i}) + N_0 * Gen_0}{N_{total}}$$

In this equation, $Gen_{t+1}$ corresponds to the effective number of generations of selection that individuals will be exposed to at the next generation. $N_t$, $N_{t-i}$, and $N_0$ correspond to the number of individuals transferred from the current generation of selection ($Gen_t$), each of the four backup boxes (from $Gen_{t-1}$ to $Gen_{t-4}$), and the replicate population of the allopatry regime ($Gen_0$), respectively, and $N_{total}$ corresponds to the total number of adult females transferred. $Gen_t$, $Gen_{t-i}$ and $Gen_0$ correspond, respectively, to the effective number of generations of selection that individuals of the current generation, the backup boxes (with $i$ ranging from 1 to 4), and the replicate populations of the allopatry regime (*i.e.*, not evolving with green females, so $Gen_0$ =

0) were exposed to; and $S_t$ corresponds to whether selection is acting on the current generation ($S_t$ = 1) or not (*i.e.*, relaxed selection in the absence of green-form females; $S_t$ = 0). Figure II provides this number for each green- and red-form population replicate of the sympatry regime.

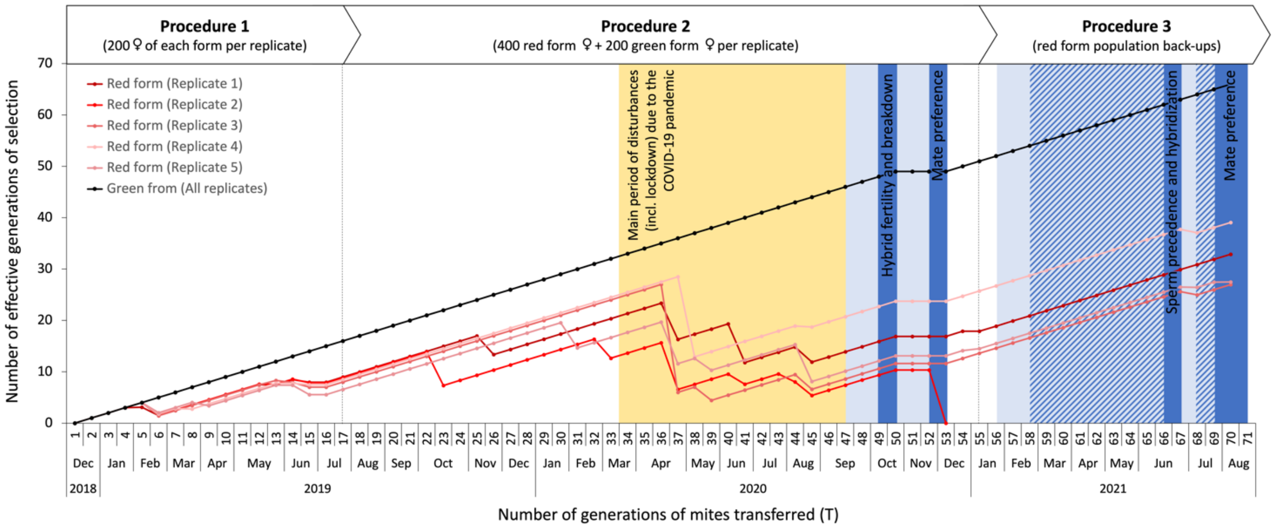

**Figure II. Number of effective generations of selection in the sympatry regime at each generation of mites transferred (T) and time at which the experiments were performed.** The number of effective generations of selection is consistent across population replicates of the green form (black line) but varies among population replicates of the red form (red lines). Drops in the number of generations of selection in the red-form population replicates were due to red females being transferred from the allopatry regime, and plateaus were due to relaxed selection (*i.e.,* when red-form populations temporarily evolved in absence of green-form individuals). The biggest issues for the maintenance of red-form mites in the sympatry regime (hence bigger drop in effective generations of selection) occurred during the first lockdown due to COVID-19 pandemic (yellow background). Individuals used in the three experiments were the offspring of females collected after different number of transfers: **T47 to assess hybrid fertility and hybrid breakdown**; **T50 and T67-69 to assess mate choice**; **T56-64 to assess sperm precedence and hybridization**. The light blue shaded backgrounds indicate the time at which the common gardens were performed ahead of each experiment, the darker blue shaded backgrounds indicate the time at which each experiment was performed, and the dashed light/dark blue shaded backgrounds indicate the overlap between common gardens and experiments.

**Table S1. Description of all statistical models used for data analyses.** For each experiment ('Exp.' column), the 'response variable' column indicates how each variable of interest was computed prior to analyses. The "data subset" column indicates which replicates were included in each analysis, and the "sample size" column gives the corresponding number of data points (*i.e.*, number of patches). The "Fixed effects in maximal model" column gives the complete set of fixed explanatory variables initially included in the model, from which non-significant fixed variables were sequentially removed to establish a minimal model ("Fixed effects in minimal model" column). The "Random effects" column indicate the variables that were fit as random explanatory variables in the model. The "R subroutine" column indicates the procedure (model function) and error structure used for each model (b: binomial, bb: betabinomial, zibb: zero-inflated betabinomial; cp: compois). **hatched/ unhatched:** number of hatched/unhatched eggs in the brood of the focal female; **dead/adults**: total number of dead juveniles/adult offspring among hatched eggs of the focal female; **daughters/sons**: total number of daughters/sons among adult offspring of the focal female; **first ♂/second ♂:** number of daughters sired by the first/second mate of the focal female; **Fertile**: whether the focal female laid at least one egg during the oviposition period; **focal/first**: colour form of the focal/first male; **mate(s)**: colour form of the male(s) with which the focal female mated; **status**: whether the focal female was mated with a single or two males; **regime**: evolution regime from which the focal individual originated; **gentime**: the generation (transfer) from which the focal individual originated; **F0 cross**: type of cross that produced the focal F1 female (Green♀ x Green ♂; Red♀ x Red ♂; Green♀ x Red ♂; or Red♀ x Green ♂); **sex-ratio**: proportion of daughters among adult offspring of the focal female; **day**: number of days after which the number of eggs was counted (either 'day 3' or 'day 5'). **block**: the experimental block in which each experimental replicate was tested; **session**: the session of behavioural observation (one session was performed per day) in which each experimental replicate was tested; **replicate**: the population replicate the females belonged to; **IDfocal**: identity of the focal female.

| Exp. | Variable of interest | Response variable | Data subset | Sample size | Model No. | Fixed effects in maximal model | Fixed effects in minimal model | Random effects | R subroutine |
|---|---|---|---|---|---|---|---|---|---|
| **Pre-mating** | Mating propensity | Mated (1) or not (0) | All ♂ | 979 | 1.1 | focal*regime*gentime | focal*regime | (1\|block) + (1\|session) + (1\|replicate) | glmmTMB[b] |
| | Mating preference | Mated with red (1) or green (0) females | All ♂ | 878[1] | 1.2 | focal*regime*gentime | regime*gentime | (1\|block) + (1\| session) + (1\|replicate) | glmmTMB[b] |
| | Latency to copulation | Number of seconds before copulation | All ♂ | 878[1] | 1.3 | focal*regime*gentime*choice | focal*regime*gentime*choice | (1\|block) + (1\| session) + (1\|replicate) | coxme |
| | Copulation duration | Number of seconds spent in copulation | All ♂ | 876[2] | 1.4 | focal*regime*gentime*choice | focal*regime + gentime*regime | (1\|block) + (1\| session) + (1\|replicate) | coxme |
| **Early post-mating** | F1 zygote mortality | cbind(unhatched, hatched) | Single-mated ♀ | 436[3] | 2.2.1 | mate*regime | mate*regime | (1\|block) + (1\|replicate) | glmmTMB[bb] |
| | | | | | 2.2.2 | mate*regime + sex-ratio | sex-ratio | (1\|block) + (1\|replicate) | glmmTMB[bb] |
| | F1 juvenile mortality | cbind(dead, adults) | Single-mated ♀ | 435[4] | 2.3.1 | mate*regime | mate*regime | (1\|block) + (1\|replicate) | glmmTMB[bb] |
| | | | | | 2.3.2 | mate*regime + sex-ratio | sex-ratio | (1\|block) + (1\|replicate) | glmmTMB[bb] |
| | F1 daughters of the 1st ♂ | cbind(first ♂, second ♂) | Double-mated ♀ | 330[5] | 2.4 | mates*regime | mates | (1\|block) + (1\|replicate) | glmmTMB[bb] |
| | F1 sex-ratio | cbind(daughters, sons) | All ♀ | 822[4] | 2.5 | first*status*regime | first*status + regime | (1\|block) + (1\|replicate) | glmmTMB[zibb] |
| **Late post-zygotic** | Proportion of fertile F1 females | Fertile (1) or not (0)[6] after 3 days oviposition | All ♀ | 959 | 3.1.1 | F0 cross*regime | F0 cross | (1\|block) + (1\|replicate) | glmmTMB[b] |
| | | Fertile (1) or not (0)[6] after 3 or 5 days oviposition | Hybrid ♀ | 960[7] | 3.1.2 | F0 cross*regime*day | 1 | (1\|block) + (1\|replicate) + (1\|IDfocal) | glmmTMB[b] |
| | F1 females oviposition | Number of F2 eggs laid over 3 days | All ♀ | 508[3] | 3.2.1 | F0 cross*regime | F0 cross + regime | (1\|block) + (1\|replicate) | glmmTMB[cp] |
| | | Number of F2 eggs laid over 3 or 5 days | Hybrid ♀ | 90[3] | 3.2.2 | F0 cross*regime*day | regime*day | (1\|block) + (1\|replicate) + (1\|IDfocal) | glmmTMB[cp] |
| | F2 zygote mortality | cbind(unhatched, hatched)[8] | Green & Red ♀ | 465 | 3.3.1 | F0 cross*regime | 1 | (1\|block) + (1\|replicate) | glmmTMB[bb] |
| | | cbind(unhatched, hatched)[8] | Hybrid ♀ | 43 | 3.3.2 | F0 cross*regime | F0 cross*regime | (1\|block) + (1\|replicate) | glmmTMB[b] |

[1] Excludes replicates for which information was missing.
[2] Excludes two additional outliers for which copulation duration lasted ca. 15 and 18 minutes, whereas all others lasted for less than 10 minutes.
[3] Excludes replicates where no egg was laid.
[4] Excludes replicates where no egg hatched.
[5] Excludes replicates where no daughters were produced.
[6] Corresponds to females that laid at least one egg (1) *versus* females that do not lay a single egg (0).
[7] Corresponds to twice the number of tested F1 hybrid females, as female fertility was assessed both after 3 and 5 days of oviposition.
[8] Corresponds to the hatching rate of eggs laid during either 3 or 5 days (in models 3.3.1 and 3.3.2, respectively).

**Box S3. Computing the strength of reproductive isolation barriers.**

We followed a method previously adapted from [1] by [2], to estimate the strength of each reproductive isolation barrier ($RI_n$) occurring at a stage $n$ in the life history of a given organism, their absolute contribution to reducing gene flow between populations ($C_n$), as well as total reproductive isolation ($T$). The formulae used to estimate the strength of each reproductive barriers are depicted on Figure III.

**Premating isolation ($\boldsymbol{RI_1}$)**
As in our previous study [3], we used an index adapted from [4-5] by [6], which represents the degree to which a population is isolated from another due to mating preferences. This index ranges from -1 to 1, positive values indicating assortative mating, a 0 value indicating random mating, and negative values indicating disassortative mating.

**Homotypic sperm precedence ($\boldsymbol{RI_2}$)**
As in [7], we used an index with the same formulation as for the previous reproductive barrier and others below, such that this index represents the degree to which homotypic (/conspecific) sperm arriving in second position in a female's reproductive track, outcompetes heterotypic (/heterospecific) sperm despite first male sperm precedence. Because this term computes the proportion of daughters sired by a homotypic second mate relative to that of a heterotypic first mate of a double mated female, it accounts for the fact that first male sperm precedence may be leaking in this system [8] (i.e., there might be some degree of shared paternity even when the sperm of a heterotypic male arrives after that of a homotypic male). As before, this index ranges between -1 and 1, with positive values indicating homotypic sperm precedence, 0 values indicating complete first male sperm precedence, and negative values indicating heterotypic sperm precedence.

**Fertilization failure ($\boldsymbol{RI_3}$)**
As in our previous study [3], we used an index adapted from [9-10] by [11]. Also referred to as "MD-type incompatibility" (MD for « Male Development » [11-15]), it corresponds to the overproduction of F1 males by haplodiploid females single-mated with an incompatible (heterotypic/heterospecific) male, as compared to females single-mated with a compatible (homotypic/conspecific) male. Indeed, an overproduction of male offspring generally indicates fertilization failure in arrhenotokous haplodiploids, because fertilized eggs develop as (diploid) females, whereas unfertilized eggs develop as (haploid) males. Although complete haploidization of fertilized eggs (e.g., due to paternal genome elimination following fertilization) could be an alternative [16-17], spider mite males are naturally produced from unfertilized eggs (i.e., true arrhenotokous parthenogenesis [18]), not from the elimination of the paternal genome in fertilized eggs (i.e., pseudoarrhenotoky [19]). Fertilization failure resulting from a defect at any of the successive stages of the reproductive process in the female reproductive tract (e.g., reduction in sperm transfer/storage, sperm ejection/dumping, reduced sperm activation or attraction to the egg, and sperm-egg incompatibility) is thus a more likely explanation for this type of incompatibility between populations [20-21]. This index ranges from 0 to 1, with positive values indicating higher fertilisation failure in heterotypic crosses than in homotypic crosses, and 0 values indicating no fertilization issue.

**Hybrid inviability ($\boldsymbol{RI_4}$)**
As for the previous reproductive isolation barrier $RI_3$, we used an index adapted from [9-10] by [11], which is also referred to as FM-type incompatibility (FM for « Female Mortality » [11-15]) because it corresponds to the proportion of unhatched eggs (hence embryonic mortality) versus female offspring in the brood of females mated with an incompatible (heterotypic/heterospecific) male, as compared to females mated with a compatible (homotypic/conspecific) male. Because in arrhenotokous haplodiploids only females among F1 offspring can be hybrids (because males are produced from unfertilized eggs; see above), this index is a good proxy for F1 hybrid inviability. As for the former index, this one ranges from 0 to 1, with positive values indicating higher mortality in heterotypic crosses than in homotypic crosses, and 0 values indicating no viability issue.

**Hybrid sterility ($RI_5$)**

We used the same formulation as for $RI_1$ and $RI_2$, such that this index represents the degree to which the fecundity of F1 hybrid females differs from that of F1 'parental type' females (from the same maternal line). Hence, this index not only accounts for the proportion of fully sterile hybrids (as in our previous study [3]), but it also encompasses the fecundity reduction in 'partially' fertile hybrids. As for $RI_1$ and $RI_2$, it ranges from -1 to 1, with positive values indicating hybrid sterility, and negative values indicating heterosis (i.e., greater fertility of the hybrids than both parents).

### Hybrid breakdown ($RI_6$)

As for $RI_2$ and $RI_3$, we used the formulation adapted by [11], to estimate the "Zygote Mortality" (ZM) in the brood of F1 hybrid females (which laid at least 1 egg) relative to that of F1 'parental type' females from the same maternal line, thereby accounting for background mortality. This index thus ranges from 0 to 1, with positive values indicating higher mortality in the brood of hybrid females, and 0 values indicating no viability issue.

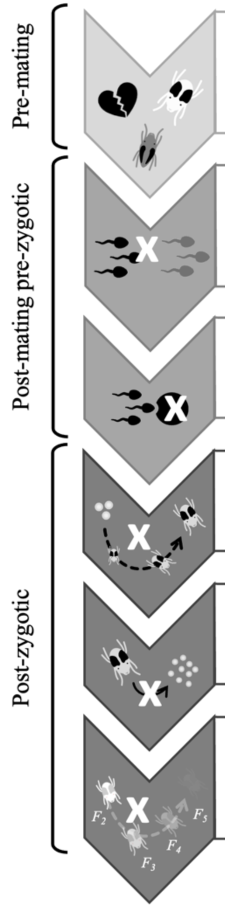

| Life stage | Reproductive barriers | Index | Measured variables |
|---|---|---|---|
| Pre-mating | **Premating isolation** *Potential mates choose not to mate due to heterospecific recognition or genitalia incompatibility.* | $RI_1 = \frac{N_{xx} - N_{xy}}{N_{xx} + N_{xy}}$ | $N_{xx}$ is the number of homotypic copulations and $N_{xy}$ is the number of heterotypic copulations. |
| Post-mating pre-zygotic | **Conspecific/homotypic sperm precedence** *Male gametes are transferred but not used when conspecific/homotypic male gametes are also present (cryptic choice).* | $RI_2 = \frac{S_{xy} - S_{yx}}{S_{xy} + S_{yx}}$ | $S_{xy}$ is the proportion of $x$ daughters sired by a $x$ male mated before a $y$ male, and $S_{yx}$ is the proportion of $x$ daughters sired by a $y$ male mated before a $x$ male. |
| | **Fertilization failure** *Male gametes are transferred, but fertilization fails.* | $RI_3 = \frac{MD_{xy} - \widehat{MD_{xx}}}{1 - \widehat{MD_{xx}}}$ | $MD_{xy}$ is the proportion of F1 males among eggs (*i.e.* male development) produced in heterotypic crosses, and $\widehat{MD_{xx}}$ is the average of the same proportion in F0 homotypic crosses. |
| Post-zygotic | **Hybrid inviability** *F1 hybrids die during embryonic or juvenile development, or as adults.* | $RI_4 = \frac{FM_{xy} - \widehat{FM_{xx}}}{1 - \widehat{FM_{xx}}}$ | $FM_{xy}$ is the proportion of F1 unhatched eggs among females (*i.e.* female mortality) produced in heterotypic crosses, and $\widehat{FM_{xx}}$ is the average of the same proportion in homotypic crosses. |
| | **Hybrid sterility** *F1 hybrids survive but are sterile.* | $RI_5 = \frac{E_x - E_h}{E_x + E_h}$ | $E_h$ is the number of eggs laid by F1 hybrids, and $E_x$ is the number of eggs laid by F1 'parental type' females. |
| | **Hybrid breakdown** *Backcrosses and/or hybrids from subsequent generations are inviable or sterile.* | $RI_6 = \frac{ZM_{xy} - \widehat{ZM_{xx}}}{1 - \widehat{ZM_{xx}}}$ | $ZM_{xy}$ is the proportion of F2 unhatched eggs among eggs (*i.e.* zygote mortality) produced in heterotypic crosses and $\widehat{ZM_{xx}}$ is the average of the same proportion in homotypic crosses. |

**Contribution of each reproductive barrier to reducing gene flow at a stage $n$ in life history:** $C_n = RI_n\,(1 - \sum_{i=1}^{n-1} C_i)$

**Total reproductive isolation for $m$ reproductive barriers:** $T = \sum_{i=1}^{m} C_i$

**Figure III. Summary of the reproductive barriers occurring at different stages in life history of spider mites, the index used to estimate the strength of each barrier (*RIₙ*) from experimentally measured variables, and the formulae used to compute their relative contribution to reducing gene flow between populations (*Cₙ*) as well as total reproductive isolation (T).**

**Table S2. Results of the post-hoc comparisons performed between data obtained for males of different colour forms and evolution regimes in the experiment testing for pre-mating isolation.** Grey cells highlight statistically significant comparisons at the 5% level. Holm corrections were used to account for multiple testing. T50 and T67-69 correspond, respectively, to 50 and 67-69 transferred generations.

| Comparison | estimate | SE | *z* ratio | p-value |
|---|---|---|---|---|
| **(a) Mating propensity – Model 1.1 (Fig. 3A)** | | | | |
| Green ♂ - allopatry *vs.* sympatry | -0.624 | 0.313 | -1.995 | 0.0460 |
| Red ♂ - allopatry *vs.* sympatry | 0.228 | 0.315 | 0.723 | 0.4697 |
| **(b) Mating preference – Model 1.2 (Fig. 3B)** | | | | |
| Allopatry - T50 *vs.* T67-69 | -0.0835 | 0.241 | -0.346 | 0.7295 |
| Sympatry - T50 *vs.* T67-69 | 0.6547 | 0.275 | 2.383 | 0.0172 |
| **(c) Latency to copulation – Model 1.3 (Fig. 3C)** | | | | |
| ♂ mated with green ♀ after T50 | | | | |
| Allopatry - green *vs.* red ♂ | 0.5483 | 0.354 | 1.548 | 0.6079 |
| Sympatry - green *vs.* red ♂ | -0.3889 | 0.460 | -0.846 | 1.0000 |
| Green ♂ - allopatry *vs.* sympatry | 0.7552 | 0.431 | 1.753 | 0.4780 |
| Red ♂ - allopatry *vs.* sympatry | -0.1820 | 0.390 | -0.467 | 1.0000 |
| Allopatry green ♂ *vs.* sympatry red ♂ | 0.3663 | 0.375 | 0.976 | 1.0000 |
| Allopatry red ♂ *vs.* sympatry green ♂ | 0.2068 | 0.443 | 0.467 | 1.0000 |
| ♂ mated with red ♀ after T50 | | | | |
| Allopatry - green *vs.* red ♂ | 0.1829 | 0.218 | 0.840 | 1.0000 |
| Sympatry - green *vs.* red ♂ | 0.2761 | 0.208 | 1.329 | 0.9187 |
| Green ♂ - allopatry *vs.* sympatry | 0.0390 | 0.216 | 0.181 | 1.0000 |
| Red ♂ - allopatry *vs.* sympatry | 0.1322 | 0.209 | 0.631 | 1.0000 |
| Allopatry green ♂ *vs.* sympatry red ♂ | 0.3151 | 0.220 | 1.431 | 0.9149 |
| Allopatry red ♂ *vs.* sympatry green ♂ | -0.1438 | 0.204 | -0.705 | 1.0000 |
| ♂ mated with green ♀ after T67-69 | | | | |
| Allopatry - green *vs.* red ♂ | -0.1347 | 0.224 | -0.601 | 0.7394 |
| Sympatry - green *vs.* red ♂ | 0.6067 | 0.212 | 2.868 | 0.0248 |
| Green ♂ - allopatry *vs.* sympatry | -0.4187 | 0.228 | -1.838 | 0.3302 |
| Red ♂ - allopatry *vs.* sympatry | 0.3227 | 0.208 | 1.550 | 0.4844 |
| Allopatry green ♂ *vs.* sympatry red ♂ | 0.1881 | 0.210 | 0.897 | 0.7394 |
| Allopatry red ♂ *vs.* sympatry green ♂ | -0.2840 | 0.227 | -1.253 | 0.6309 |
| ♂ mated with red ♀ after T67-69 | | | | |
| Allopatry - green *vs.* red ♂ | 0.0806 | 0.131 | 0.614 | 1.0000 |
| Sympatry - green *vs.* red ♂ | 0.0821 | 0.134 | 0.615 | 1.0000 |
| Green ♂ - allopatry *vs.* sympatry | 0.0436 | 0.130 | 0.334 | 1.0000 |
| Red ♂ - allopatry *vs.* sympatry | 0.0451 | 0.135 | 0.333 | 1.0000 |
| Allopatry green ♂ *vs.* sympatry red ♂ | 0.1257 | 0.135 | 0.928 | 1.0000 |
| Allopatry red ♂ *vs.* sympatry green ♂ | -0.0370 | 0.130 | -0.285 | 1.0000 |
| **(d) Copulation duration – Model 1.4 (Fig. 3D)** | | | | |
| ♂ mated with any ♀ after T50 | | | | |
| Allopatry - green *vs.* red ♂ | -0.547 | 0.0981 | -5.580 | <.0001 |
| Sympatry - green *vs.* red ♂ | -0.908 | 0.0988 | -9.191 | <.0001 |
| Green ♂ - allopatry *vs.* sympatry | -0.100 | 0.1540 | -0.648 | 0.5167 |
| Red ♂ - allopatry *vs.* sympatry | -0.461 | 0.1550 | -2.969 | 0.0090 |
| Allopatry green ♂ *vs.* sympatry red ♂ | -1.008 | 0.1590 | -6.350 | <.0001 |
| Allopatry red ♂ *vs.* sympatry green ♂ | -0.447 | 0.1530 | -2.928 | 0.0090 |
| ♂ mated with any ♀ after T67-69 | | | | |
| Allopatry - green *vs.* red ♂ | -0.547 | 0.0981 | -5.580 | <.0001 |
| Sympatry - green *vs.* red ♂ | -0.908 | 0.0988 | -9.191 | <.0001 |
| Green ♂ - allopatry *vs.* sympatry | 0.224 | 0.1130 | 1.978 | 0.0959 |
| Red ♂ - allopatry *vs.* sympatry | -0.137 | 0.1120 | -1.216 | 0.2240 |
| Allopatry green ♂ *vs.* sympatry red ♂ | -0.684 | 0.1140 | -6.019 | <.0001 |
| Allopatry red ♂ *vs.* sympatry green ♂ | -0.771 | 0.1150 | -6.720 | <.0001 |

**Table S3. Mating propensity and mate choice observed in each replicate population in the experiment testing for pre-mating isolation.** For each evolution regime, tested generation (number of experimental evolution transfers), colour form of focal males ('Focal ♂' column), and replicate population ('Rep.' column), the table gives the total number of tested males ($N_{total}$), the total number and proportion (± s.e.) of males mated with one of the proposed females ('$N_{mated}$' and 'Mating propensity', respectively), and the total number of males mated with each of the two possible mates ('$N_{mated}$ red ♀' and '$N_{mated}$ green ♀'), and the mean proportion (± s.e.) of males mated with a red female ('Preference for red ♀').

| Regime | Generation | Focal ♂ | Rep. | $N_{total}$ | $N_{mated}$ | Mating propensity (%) | $N_{mated\ red}$ ♀ | $N_{mated\ green}$ ♀ | Preference for red ♀ (%) |
|---|---|---|---|---|---|---|---|---|---|
| Allopatry | T50 | Green ♂ | 1 | 16 | 14 | 87.50 ± 8.27 | 8 | 6 | 57.14 ± 13.23 |
| | | | 3 | 16 | 16 | 100.0 ± 0.00 | 9 | 7 | 56.25 ± 12.40 |
| | | | 4 | 16 | 15 | 93.75 ± 6.05 | 12 | 3 | 80.00 ± 10.33 |
| | | | 5 | 16 | 13 | 81.25 ± 9.76 | 11 | 2 | 84.62 ± 10.01 |
| | | Red ♂ | 1 | 16 | 14 | 87.50 ± 8.27 | 10 | 4 | 71.43 ± 12.07 |
| | | | 3 | 16 | 16 | 100.0 ± 0.00 | 13 | 3 | 81.25 ± 9.76 |
| | | | 4 | 16 | 16 | 100.0 ± 0.00 | 12 | 4 | 75.00 ± 10.83 |
| | | | 5 | 16 | 16 | 100.0 ± 0.00 | 12 | 4 | 75.00 ± 10.83 |
| | T67-69 | Green ♂ | 1 | 47 | 42 | 89.36 ± 4.50 | 37 | 5 | 88.10 ± 5.00 |
| | | | 3 | 45 | 38 | 84.44 ± 5.40 | 27 | 11 | 71.05 ± 7.36 |
| | | | 4 | 45 | 38 | 84.44 ± 5.40 | 25 | 13 | 65.79 ± 7.70 |
| | | | 5 | 45 | 38 | 84.44 ± 5.40 | 27 | 11 | 71.05 ± 7.36 |
| | | Red ♂ | 1 | 45 | 40 | 88.89 ± 4.68 | 29 | 11 | 72.50 ± 7.06 |
| | | | 3 | 45 | 38 | 84.44 ± 5.40 | 26 | 12 | 68.42 ± 7.54 |
| | | | 4 | 45 | 42 | 93.33 ± 3.72 | 31 | 11 | 73.81 ± 6.78 |
| | | | 5 | 45 | 41 | 91.11 ± 4.24 | 33 | 8 | 80.49 ± 6.19 |
| Sympatry | T50 | Green ♂ | 1 | 16 | 15 | 93.75 ± 6.05 | 11 | 4 | 73.33 ± 11.42 |
| | | | 3 | 16 | 14 | 87.50 ± 8.27 | 11 | 3 | 78.57 ± 10.97 |
| | | | 4 | 16 | 15 | 93.75 ± 6.05 | 15 | 0 | 100.0 ± 0.00 |
| | | | 5 | 14 | 14 | 100.0 ± 0.00 | 13 | 1 | 92.86 ± 6.88 |
| | | Red ♂ | 1 | 16 | 14 | 87.50 ± 8.27 | 12 | 2 | 85.71 ± 9.35 |
| | | | 3 | 16 | 16 | 100.0 ± 0.00 | 13 | 3 | 81.25 ± 9.76 |
| | | | 4 | 15 | 14 | 93.33 ± 6.44 | 10 | 4 | 71.43 ± 12.07 |
| | | | 5 | 14 | 13 | 92.86 ± 6.88 | 10 | 3 | 76.92 ± 11.69 |
| | T67-69 | Green ♂ | 1 | 45 | 43 | 95.56 ± 3.07 | 35 | 7 | 83.33 ± 5.75 |
| | | | 3 | 45 | 39 | 86.67 ± 5.07 | 26 | 13 | 66.67 ± 7.55 |
| | | | 4 | 44 | 39 | 88.64 ± 4.78 | 30 | 9 | 76.92 ± 6.75 |
| | | | 5 | 45 | 43 | 95.56 ± 3.07 | 33 | 10 | 76.74 ± 6.44 |
| | | Red ♂ | 1 | 46 | 39 | 84.78 ± 5.30 | 27 | 12 | 69.23 ± 7.39 |
| | | | 3 | 45 | 40 | 88.89 ± 4.68 | 22 | 18 | 55.00 ± 7.87 |
| | | | 4 | 51 | 44 | 86.27 ± 4.82 | 31 | 11 | 73.81 ± 6.78 |
| | | | 5 | 45 | 42 | 93.33 ± 3.72 | 28 | 14 | 66.67 ± 7.27 |

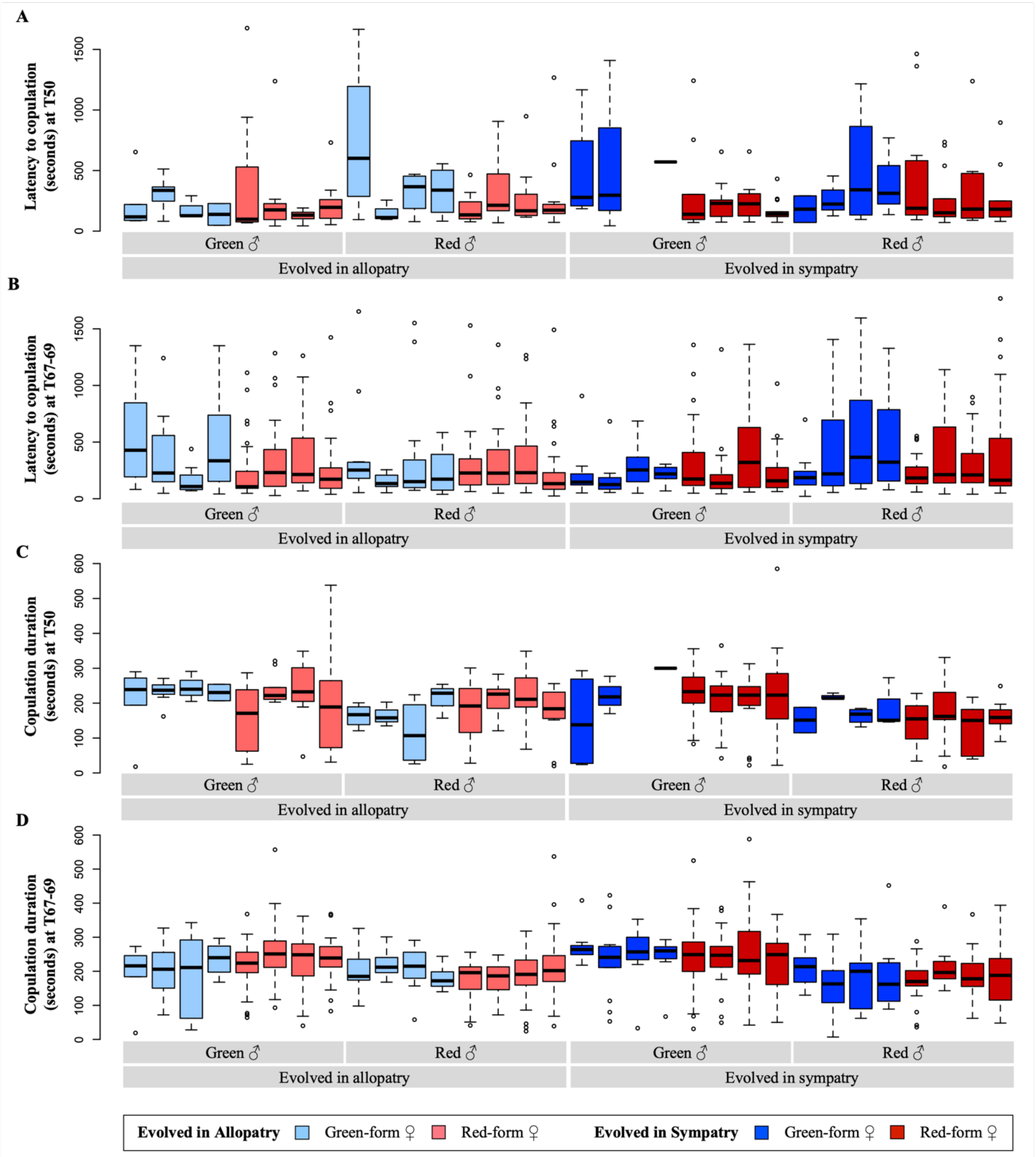

**Figure S1. Latency to copulation and copulation duration with red- and green-form females for red- and green-form males evolved in each replicate population of each evolution regime.** Boxplots show, for each of the four replicate populations (Rep. 1, 3, 4 and 5 from left to right) in each evolution regime, the median and quartiles of latency to copulation (A, B) and copulation duration (C, D) measured (in seconds) after 50 (A, C) and 67-69 (B, D) transfers of experimental evolution for each male forms. Lighter and darker colour tones represent individuals evolved in the allopatry and sympatry regimes, respectively. Blue: green-form females; Red: red-form females.

**Table S4. Results of the post-hoc comparisons performed for F1 zygote mortality (a), F1 juvenile mortality (b), and sex ratio (c) data obtained for green-form females from different evolution regimes and mated with different males in the experiment testing for early post-mating isolation.** Light grey cells highlight statistically significant comparisons at the 5% level. Holm corrections were used to account for multiple testing.

| | **estimate** | **SE** | ***z* ratio** | **p-value** |
|---|---|---|---|---|
| **(a) F1 zygote mortality – Model 2.2.1 (without sex ratio as covariate; Fig. 4A)** | | | | |
| Allopatry - Green ♂ *vs.* Red ♂ | -0.163 | 0.157 | -1.035 | 0.6018 |
| Green ♂ - Allopatry *vs.* Sympatry | 0.337 | 0.168 | 2.011 | 0.1774 |
| Green ♂ Allopatry *vs.* Red ♂ Sympatry | -0.292 | 0.158 | -1.849 | 0.1934 |
| Red ♂ Allopatry *vs.* Green ♂ Sympatry | 0.500 | 0.168 | 2.985 | 0.0142 |
| Red ♂ - Allopatry *vs.* Sympatry | -0.130 | 0.157 | -0.824 | 0.6018 |
| Sympatry - Green ♂ *vs.* Red ♂ | -0.630 | 0.168 | -3.747 | 0.0011 |
| **(b) F1 juvenile mortality – Model 2.3.1 (without sex ratio as covariate; Fig. 4B)** | | | | |
| Allopatry - Green ♂ *vs.* Red ♂ | 0.3205 | 0.123 | 2.601 | 0.0372 |
| Green ♂ - Allopatry *vs.* Sympatry | -0.3711 | 0.154 | -2.412 | 0.0476 |
| Green ♂ Allopatry *vs.* Red ♂ Sympatry | 0.3538 | 0.167 | 2.122 | 0.0677 |
| Red ♂ Allopatry *vs.* Green ♂ Sympatry | -0.6916 | 0.161 | -4.299 | 0.0001 |
| Red ♂ - Allopatry *vs.* Sympatry | 0.0333 | 0.173 | 0.193 | 0.8472 |
| Sympatry - Green ♂ *vs.* Red ♂ | 0.7248 | 0.124 | 5.839 | <.0001 |
| **(c) F1 sex ratio – Model 2.5 (Fig. 4D)** | | | | |
| single-mated females - green *vs.* red ♂ | 1.8484 | 0.126 | 14.617 | <.0001 |
| double-mated females - green *vs.* red ♂ | 1.4005 | 0.130 | 10.784 | <.0001 |
| Green ♂ - single- vs. double-mated females | -0.0401 | 0.115 | -0.349 | 0.7274 |
| Red ♂ - single- vs. double-mated females | -0.4880 | 0.135 | -3.619 | 0.0006 |

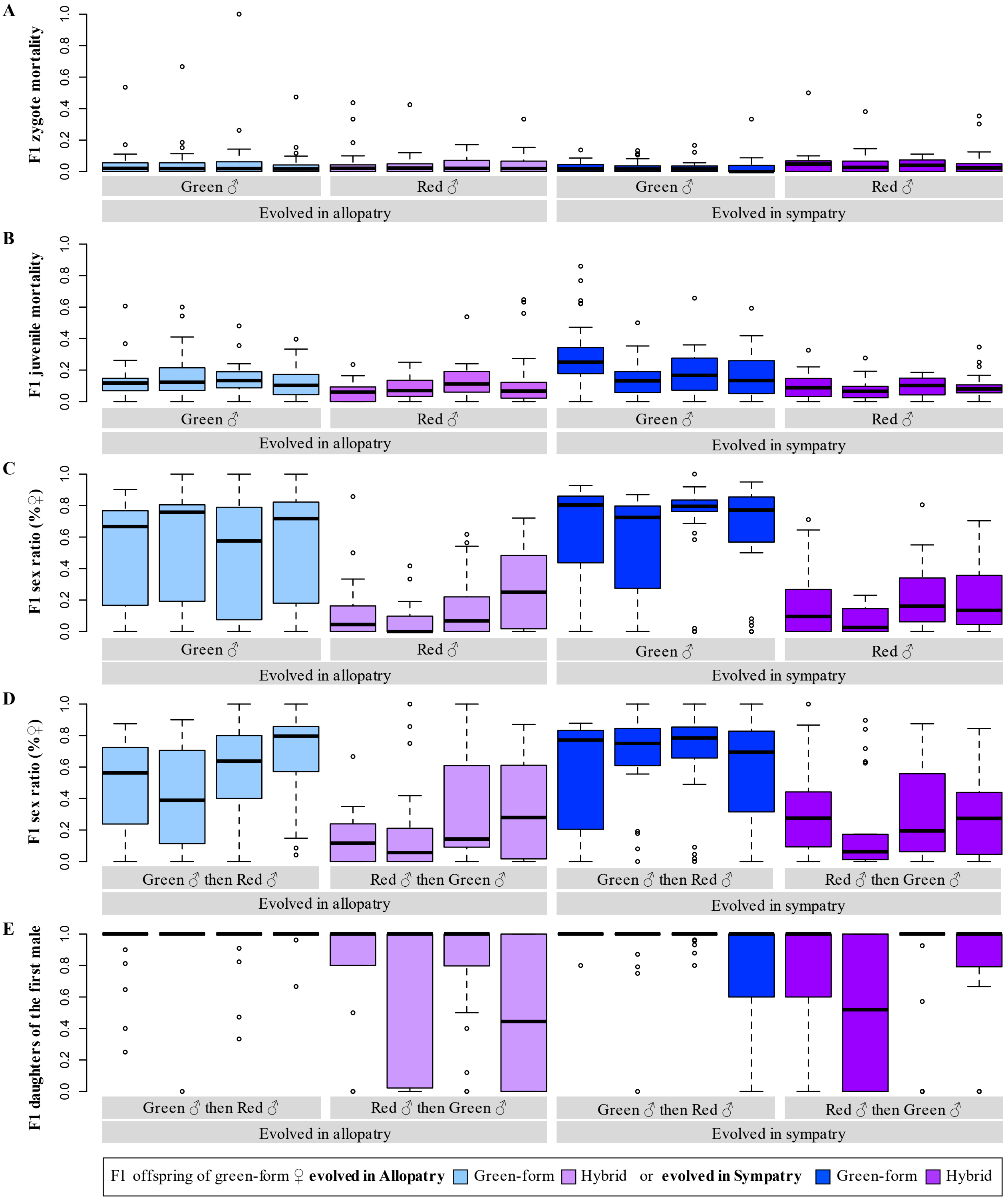

**Figure S2. F1 zygote and juvenile mortality, sex ratio, and sperm precedence in single and double mated green-form females in each replicate population of each evolution regime.** Boxplots show, for each of the four replicate populations (Rep. 1, 3, 4 and 5 from left to right) in each evolution regime, the median and quartiles of the proportion of (A) unhatched eggs, (B) dead juveniles, (C) daughters among adult offspring in the brood of single mated green-form females, (D) daughters among adult offspring in the brood of double mated green females, and among the latter, (E) daughters sired by the first mate only. In all panels, the x axis indicates the evolution regime and the colour form of the male(s) mated with the females. Lighter and darker colour tones represent crosses from the allopatry and from the sympatry regime, respectively. Blue: homotypic crosses (green-form offspring); Purple: heterotypic crosses (hybrid daughters in panel E, or offspring incl. hybrid daughters in all other panels).

**Table S5. Proportion of F1 fertile females and number of eggs laid in each replicate population after 3 or 5 days of oviposition.** The table gives, for each replicate population ('Rep.') of each evolution regime, the total number and mean (± s.e.) proportion of focal F1 females, resulting from different ♀ x ♂ F0 cross (G: green; R: red), that laid at least one egg ('% fertile' column), and average number (± s.e.) of eggs laid ('No. eggs' column) by these oviositing females (i.e., excluding fully sterile females) after 3 days of oviposition, as well as after 5 days oviposition for hybrid females . $N_{total}$: total number of tested focal F1 females. $N_{fertile}$: total number of focal females that laid at lead one egg.

| | | 3 days oviposition | | | | 5 days oviposition | | | |
|---|---|---|---|---|---|---|---|---|---|
| Focal F1♀ | Rep. | $N_{total}$ | $N_{fertile}$ | % fertile | No. eggs | $N_{total}$ | $N_{fertile}$ | % fertile | No. eggs |
| Green♀ (GxG F0 cross) evolved in **allopatry** | 1 | 24 | 22 | 91.67 ± 5.64 | 25.18 ± 1.39 | - | - | - | - |
| | 2 | 24 | 22 | 91.67 ± 5.64 | 26.05 ± 1.6 | - | - | - | - |
| | 3 | 24 | 24 | 100 ± 0.00 | 26.54 ± 1.46 | - | - | - | - |
| | 4 | 24 | 24 | 100 ± 0.00 | 24.29 ± 1.47 | - | - | - | - |
| | 5 | 24 | 22 | 91.67 ± 5.64 | 26.64 ± 1.02 | - | - | - | - |
| Hybrid♀ (GxR F0 cross) evolved in **allopatry** | 1 | 24 | 5 | 20.83 ± 8.29 | 2.40 ± 0.93 | 24 | 5 | 20.83 ± 8.29 | 2.4 ± 0.93 |
| | 2 | 24 | 3 | 12.50 ± 6.75 | 3.33 ± 0.67 | 24 | 3 | 12.50 ± 6.75 | 3.67 ± 0.33 |
| | 3 | 24 | 2 | 8.33 ± 5.64 | 8.00 ± 1.00 | 24 | 3 | 12.50 ± 6.75 | 8 ± 1 |
| | 4 | 24 | 0 | 0.00 ± 0.00 | - | 24 | 0 | 0.00 ± 0.00 | - |
| | 5 | 24 | 0 | 0.00 ± 0.00 | - | 24 | 0 | 0.00 ± 0.00 | - |
| Hybrid♀ (RxG F0 cross) evolved in **allopatry** | 1 | 24 | 3 | 12.50 ± 6.75 | 5.33 ± 2.96 | 24 | 4 | 16.67 ± 7.61 | 5.67 ± 3.28 |
| | 2 | 24 | 3 | 12.50 ± 6.75 | 9 ± 7 | 24 | 3 | 12.50 ± 6.75 | 9 ± 7 |
| | 3 | 24 | 3 | 12.50 ± 6.75 | 5.67 ± 2.4 | 24 | 3 | 12.50 ± 6.75 | 6 ± 2.65 |
| | 4 | 24 | 3 | 12.50 ± 6.75 | 11.67 ± 3.67 | 24 | 4 | 16.67 ± 7.61 | 15 ± 5.57 |
| | 5 | 24 | 3 | 12.50 ± 6.75 | 6 ± 2.52 | 24 | 3 | 12.50 ± 6.75 | 6 ± 2.52 |
| Red♀ (RxR F0 cross) evolved in **allopatry** | 1 | 24 | 24 | 100 ± 0.00 | 23.58 ± 1.31 | - | - | - | - |
| | 2 | 24 | 24 | 100 ± 0.00 | 25.29 ± 0.99 | - | - | - | - |
| | 3 | 24 | 24 | 100 ± 0.00 | 25.29 ± 1.13 | - | - | - | - |
| | 4 | 24 | 24 | 100 ± 0.00 | 24.58 ± 0.77 | - | - | - | - |
| | 5 | 24 | 23 | 95.83 ± 4.08 | 24.17 ± 1.11 | - | - | - | - |
| Green♀ (GxG F0 cross) evolved in **sympatry** | 1 | 24 | 24 | 100 ± 0.00 | 19.67 ± 1.39 | - | - | - | - |
| | 2 | 23 | 21 | 91.30 ± 5.88 | 20.52 ± 1.38 | - | - | - | - |
| | 3 | 24 | 23 | 95.83 ± 4.08 | 23.35 ± 0.72 | - | - | - | - |
| | 4 | 24 | 23 | 95.83 ± 4.08 | 20.35 ± 1.1 | - | - | - | - |
| | 5 | 24 | 24 | 100 ± 0.00 | 23.38 ± 1.2 | - | - | - | - |
| Hybrid♀ (GxR F0 cross) evolved in **sympatry** | 1 | 24 | 1 | 4.17 ± 4.08 | 4 ± NA | 24 | 1 | 4.17 ± 4.08 | 4 ± NA |
| | 2 | 24 | 2 | 8.33 ± 5.64 | 3.5 ± 0.5 | 24 | 2 | 8.33 ± 5.64 | 3.5 ± 0.5 |
| | 3 | 24 | 0 | 0.00 ± 0.00 | - | 24 | 0 | 0.00 ± 0.00 | - |
| | 4 | 24 | 1 | 4.17 ± 4.08 | 13 ± NA | 24 | 1 | 4.17 ± 4.08 | 14 ± NA |
| | 5 | 24 | 4 | 16.67 ± 7.61 | 1.75 ± 0.48 | 24 | 4 | 16.67 ± 7.61 | 5.25 ± 3.92 |
| Hybrid♀ (RxG F0 cross) evolved in **sympatry** | 1 | 24 | 3 | 12.50 ± 6.75 | 19.67 ± 6.89 | 24 | 3 | 12.50 ± 6.75 | 30.33 ± 12.33 |
| | 2 | 24 | 2 | 8.33 ± 5.64 | 4 ± 2 | 24 | 2 | 8.33 ± 5.64 | 4 ± 2 |
| | 3 | 24 | 2 | 8.33 ± 5.64 | 1 ± 0 | 24 | 3 | 12.50 ± 6.75 | 1 ± 0 |
| | 4 | 24 | 1 | 4.17 ± 4.08 | 2 ± NA | 24 | 1 | 4.17 ± 4.08 | 2 ± NA |
| | 5 | 24 | 2 | 8.33 ± 5.64 | 5.5 ± 0.5 | 24 | 2 | 8.33 ± 5.64 | 5.5 ± 0.5 |
| Red♀ (RxR F0 cross) evolved in **sympatry** | 1 | 24 | 24 | 100 ± 0.00 | 22.75 ± 1.46 | - | - | - | - |
| | 2 | 24 | 23 | 95.83 ± 4.08 | 21.91 ± 1.33 | - | - | - | - |
| | 3 | 24 | 23 | 95.83 ± 4.08 | 22.13 ± 1.18 | - | - | - | - |
| | 4 | 24 | 23 | 95.83 ± 4.08 | 24.7 ± 1 | - | - | - | - |
| | 5 | 24 | 24 | 100 ± 0.00 | 24.54 ± 1.67 | - | - | - | - |

**Table S6. Results of the post-hoc comparisons performed between data obtained for F1 females resulting from different types of F0 crosses and evolution regimes in the experiment testing for late post-zygotic isolation.** Light grey cells highlight statistically significant comparisons at the 5% level. Holm corrections were used to account for multiple testing. G: green-form; R: red-form.

| Comparison | estimate | SE | *z* ratio | p-value |
|---|---|---|---|---|
| **(a) Proportion of F1 ovipositing females (Fig. 5A)** | | | | |
| Between types of ♀ x ♂ F0 crosses following 3 days of oviposition – **Model 3.1.1** | | | | |
| G x G *vs.* R x R | -0.947 | 0.599 | -1.581 | 0.2279 |
| G x R *vs.* R x G | -0.361 | 0.324 | -1.115 | 0.2650 |
| (G x G and R x R) *vs.* (G x R and R x G) | 5.945 | 0.350 | 16.992 | <0.0001 |
| **(b) Number of eggs laid by F1 ovipositing females (Fig. 5B)** | | | | |
| Between types of ♀ x ♂ F0 crosses following 3 days of oviposition – **Model 3.2.1** | | | | |
| G x G *vs.* R x R | -0.0134 | 0.0277 | -0.482 | 0.6297 |
| G x G *vs.* G x R | 1.8188 | 0.1620 | 11.225 | <.0001 |
| G x G *vs.* R x G | 1.1291 | 0.1030 | 10.963 | <.0001 |
| R x R *vs.* G x R | 1.8321 | 0.1620 | 11.311 | <.0001 |
| R x R *vs.* R x G | 1.1424 | 0.1029 | 11.100 | <.0001 |
| G x R *vs.* R x G | -0.6897 | 0.1900 | -3.631 | 0.0006 |
| Between days of oviposition and evolution regimes for hybrid females, irrespective of F0 crosses– **Model 3.2.2** | | | | |
| Allopatry day 3 *vs.* Sympatry day 3 | 0.2400 | 0.3857 | 0.622 | 1.0000 |
| Allopatry day 3 *vs.* Allopatry day 5 | -0.0816 | 0.0568 | -1.436 | 0.7548 |
| Allopatry day 3 *vs.* Sympatry day 5 | -0.1187 | 0.3842 | -0.309 | 1.0000 |
| Sympatry day 3 *vs.* Allopatry day 5 | -0.3216 | 0.3850 | -0.835 | 1.0000 |
| Sympatry day 3 *vs.* Sympatry day 5 | -0.3587 | 0.0618 | -5.803 | <.0001 |
| Allopatry day 5 *vs.* Sympatry day 5 | -0.0370 | 0.3835 | -0.097 | 1.0000 |
| **(c) F2 zygote mortality (Fig. 5D)** | | | | |
| Between types of F0 crosses used to produce the F1 hybrid females and evolution regimes – **Model 3.3.2** | | | | |
| G x R Allopatry *vs.* R x G Allopatry | -1.277 | 0.683 | 1.870 | 0.1846 |
| G x R Allopatry *vs.* G x R Sympatry | 1.175 | 0.645 | -1.822 | 0.1846 |
| G x R Allopatry *vs.* R x G Sympatry | -2.252 | 0.910 | 2.474 | 0.0534 |
| R x G Allopatry *vs.* G x R Sympatry | 2.452 | 0.567 | -4.326 | 0.0001 |
| R x G Allopatry *vs.* R x G Sympatry | -0.975 | 0.883 | 1.104 | 0.2695 |
| G x R Sympatry *vs.* R x G Sympatry | -3.428 | 0.822 | 4.171 | 0.0002 |

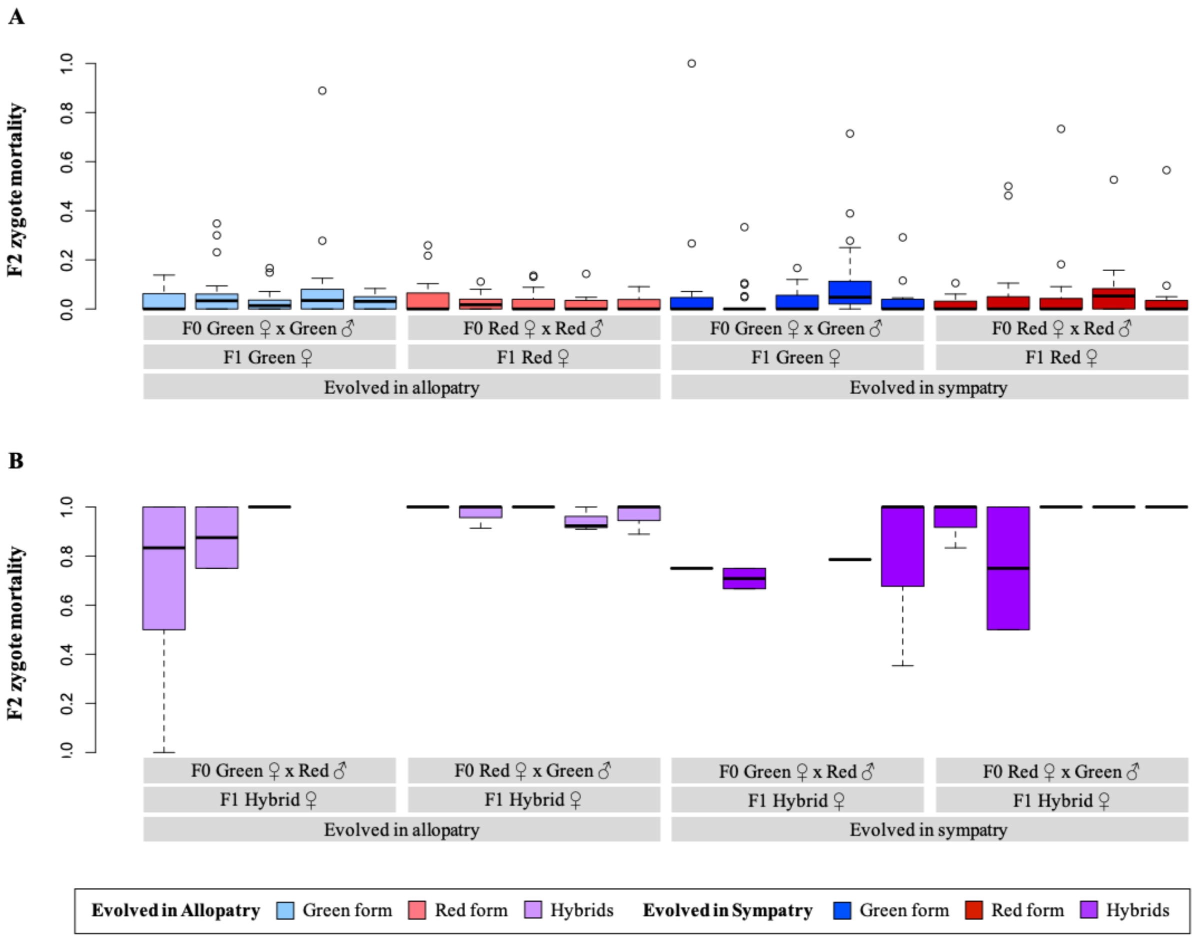

**Figure S3. Mortality of F2 eggs in each replicate population of each evolution regime.** Boxplots show the median and quartiles of the proportion of unhatched eggs in the brood of ovipositing F1 virgin females resulting from (A) homotypic and (B) hererotypic F0 crosses. Lighter and darker colour tones represent crosses from the allopatry and from the sympatry regime, respectively. Blue: homotypic F0 crosses between green females and males; Red: homotypic F0 crosses between red females and males; Purple: heterotypic F0 crosses between green females and red males or between red females and green males.

### Box S4. Sex ratio of the red- and green-form source populations in panmixia.

Sex ratio data reported here for the red-form population correspond to those obtained for the « T×T » crosses (*Wolbachia*-uninfected female × male crosses) of the population AMP (see Box S1) in a previous study [1]. Data obtained for the green-form population (*i.e.*, the population 'TOM', see Box S1) were not included in [1] because this population was not collected during the same field campaign, but it was tested simultaneously with all other populations reported in that paper. The procedure was chosen to increase potential conflicts over sex ratio, as the optimal sex ratio for female spider mites depends on the number of foundresses on a patch, being more male biased as this number increases, whereas males always benefit from female-biased sex ratio [2, 3].

#### *Experimental procedure*

The experimental procedure is fully detailed in [1]. Briefly, 10 adult virgin females were placed with 10 males with similar age on a 9 $cm^2$ bean leaf disc where they could mate in panmixia. Females could oviposit for three days, after which time the adults were removed, and the number of adult offspring of each sex was counted 15 days later. The entire experiment was done in three consecutive blocks, each including four replicates for each mite population.

#### *Statistical analyses*

Analyses were carried out using the R statistical package (v. 4.4.2; [4]). Sex ratio data were computed using the function cbind, binding together the number of females and of males counted per patch, and were analysed using a glmmTMB (glmmTMB package; [5]) with a binomial error distribution (after checking for overdispersion and zero inflation using, respectively, the testDispersion and testZeroInflation functions of the DHARMa package; [6]). The population (colour form) was fit as fixed explanatory variable, whereas the experimental block was fit as random explanatory variable. The significance of the explanatory variable was then established using a chi-squared test with the Anova function (car package; [7]).

#### *Results*

The obtained data revealed that, in panmictic conditions with 10 females per patch, the green-form source population produced, on average, 9% less females than the red-form source population ($X^2_1=14.65, p=0.0001$; Figure IV).

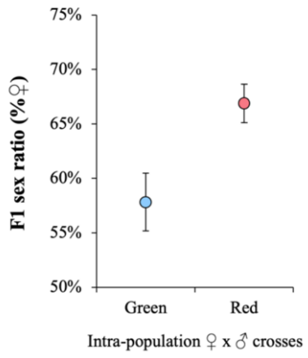

**Figure IV. Sex ratio of the red- and green-form source populations in panmictic conditions**. The offspring sex-ratio is computed here as the proportion of females among adult offspring (number of $F_1$ females /total number of $F_1$ adults).
